# N-Acetylaspartate Synthesis as a Thermodynamic Relief Mechanism for Mitochondrial Aspartate Aminotransferase

**DOI:** 10.64898/2026.06.29.735215

**Authors:** Narayanan Puthillathu, John R. Moffett, Barbara S. Slusher, Aryan M. Namboodiri

## Abstract

N-acetylaspartate (NAA) is the most abundant neuron-enriched acetylated metabolite in the mammalian brain, but its metabolic purpose remains unresolved. We developed a simplified kinetic model of mitochondrial aspartate metabolism to test whether NAA synthesis by aspartate N-acetyltransferase (ASPNAT) acts as a thermodynamic “relief valve” for mitochondrial aspartate aminotransferase (AAT) under the low-oxaloacetate (OAA) conditions expected in neuronal mitochondria. In the mitochondrial-compartment model, ASPNAT lowered steady-state mitochondrial aspartate from 141 to 105 *µ*M and increased net forward AAT flux by 30.9%. The relative AAT-relief effect was largest when OAA and aspartate-glutamate carrier 1 (AGC1/Aralar1)-mediated export were both low, whereas acetyl-CoA availability controlled the substrate-supported capacity for NAA synthesis. That places the relief effect in a narrow regime where product removal matters most. ASPNAT titration produced a graded, concentration-dependent response rather than a binary on/off switch. Energetic comparisons showed that the gain in AAT-linked support comes at a modest acetyl-CoA cost, which makes NAA synthesis easier to sustain in carbon-replete states than in carbon-poor ones. Some studies have suggested a secondary cytoplasmic site of NAA synthesis, and we therefore examined how the network response changed with a change in ASPNAT topology. Mitochondrial matrix ASPNAT increased forward AAT flux by 53.32%, whereas cytoplasmic ASPNAT decreased ASPNAT flux by 17.8%. Allowing OAA to vary preserved the positive ASPNAT-dependent relief of AAT flux, but because this simplified extension produced unrealistically low absolute fluxes, it is interpreted as a robustness check on the direction of the mechanism rather than as a prediction of physiological metabolic rates. These results suggest that ASPNAT can serve as an auxiliary matrix-aspartate sink when the AAT node is product-limited.

## 1 Introduction

NAA is the most concentrated acetylated metabolite in the mammalian brain and one of the most widely used spectroscopic indicators of neuronal health and integrity (Moffett et al., 2007; Sharma et al., 2010). Neurons synthesize NAA through aspartate N-acetyltransferase (ASPNAT; formerly known as NAT8L), which transfers an acetyl group from acetyl-CoA to aspartate (Ariyannur et al., 2010b; Goldstein, 1969; Madhavarao et al., 2003; Truckenmiller et al., 1985). ASPNAT belongs to the general control non-depressible 5 (GCN5)-related N-acetyltransferase (GNAT) family of membrane-bound acetyltransferases. NAA has been shown to provide acetate for cytoplasmic lipid synthesis, with many additional roles proposed in the literature, yet these ideas do not fully explain why neurons maintain a large, continuously synthesized NAA pool (Cambron et al., 2012; Francis et al., 2016; Mehta and Namboodiri, 1995; Moffett et al., 2013; Moffett et al., 2020b; Moffett et al., 2007). Classic biochemical studies placed NAA synthesis at a central node in mitochondrial carbon metabolism wherein the acetyl group of NAA supports brain lipid synthesis, emphasizing that NAA production supports acetate flux from mitochondria and extramitochondrial acetyl-CoA production (D’Adamo et al., 1968; D’Adamo and Yatsu, 1966; Patel and Clark, 1979, 1980, 1981). Placing NAA synthesis at the core of energy derivation pathways in the mitochondrial matrix raises the possibility it may serve an immediate mitochondrial function in addition to its better-known downstream roles in cytoplasmic lipid synthesis. NAA synthesis in the healthy adult brain is slow, but continuous (Xu et al., 2008). This continuous synthesis of NAA, even in the face of the very high NAA concentrations maintained in neurons, could provide an energetic benefit in mitochondrial metabolism beyond transport of an acetyl-CoA precursor to the cytoplasm.

Here we propose that mitochondrial ASPNAT acts as an auxiliary aspartate sink. By converting matrix aspartate to NAA, ASPNAT can pull the near-equilibrium GOT2 reaction forward when OAA is low and AGC1-mediated export is insufficient. In this model, NAA synthesis is therefore not simply a downstream biosynthetic pathway, but a mechanism that can help maintain mitochondrial carbon flux, thus facilitating neuronal energy derivation.

That possibility acquires additional support when NAA is placed in the wider context of neuronal glutamate handling. Neurons depend on astrocyte-supported glutamine recycling, transporter-mediated glutamate capture, and rapid mitochondrial transamination to connect neurotransmitter carbon with oxidative metabolism (Albrecht et al., 2007; Farooqui et al., 2008; Mates et al., 2013; McNair et al., 2020; Sonnewald and Kondziella, 2003). Within that circuit, the mitochondrial aspartate node contributes to what has often been described as a truncated neuronal tricarboxylic-acid cycle, in which glutamate-derived carbon is routed into *α*-ketoglutarate oxidation and malate-aspartate-shuttle (MAS) exchange (Gruetter, 2009; Hertz and Rodrigues, 2014; Hirrlinger and Nave, 2014; Rodrigues and Cerdán, 2007; Ryan et al., 2019). Synaptosome flux measurements further showed that AAT and the MAS interact on different kinetic scales within the same metabolic neighborhood, supporting the idea that product handling at this node can alter carbon routing (Yudkoff et al., 1994). In that setting, any pathway that changes local product pressure on mitochondrial AAT can influence more than aspartate flux alone; it can reshape how efficiently neuronal glutamate carbon couples to redox transfer and oxidative ATP production. Two major isoforms of AAT exist, GOT1 (glutamic oxaloacetic transaminase 1) is the cytoplasmic form of the enzyme, whereas GOT2 is the mitochondrial form. Our model examines the effects of NAA synthesis on GOT2, the intra-mitochondrial isoform.

One longstanding possibility is that NAA synthesis acts as a relief pathway supporting mitochondrial AAT throughput in neurons, thereby promoting glutamate oxidation and MAS flux by limiting product accumulation (Brunner et al., 2026; Madhavarao et al., 2003; Menga et al., 2021; Moffett et al., 2007; Sellinger et al., 1962; Yudkoff et al., 1994). This hypothesis is attractive because neuronal mitochondrial AAT (GOT2) operates close to thermodynamic equilibrium, while mitochondrial oxaloacetate (OAA) concentrations remain extremely low because it is continuously consumed by citrate synthase and constrained by malate dehydrogenase activity (Cooper et al., 2023; Erecinska et al., 1988; Kimmich et al., 2002; McKenna et al., 2006b; Morgunov and Srere, 1998; Thangavelu et al., 2017). Under those conditions, forward AAT flux is not expected to be controlled primarily by enzyme activity. It should instead be highly sensitive to nutrient supply and the balance between product accumulation and product removal. ASPNAT consumes mitochondrial aspartate directly, thermodynamically facilitating AAT flux and generating an acetyl-CoA precursor via a citrate synthase-independent pathway (**Figure 1**).

**Figure 1.**
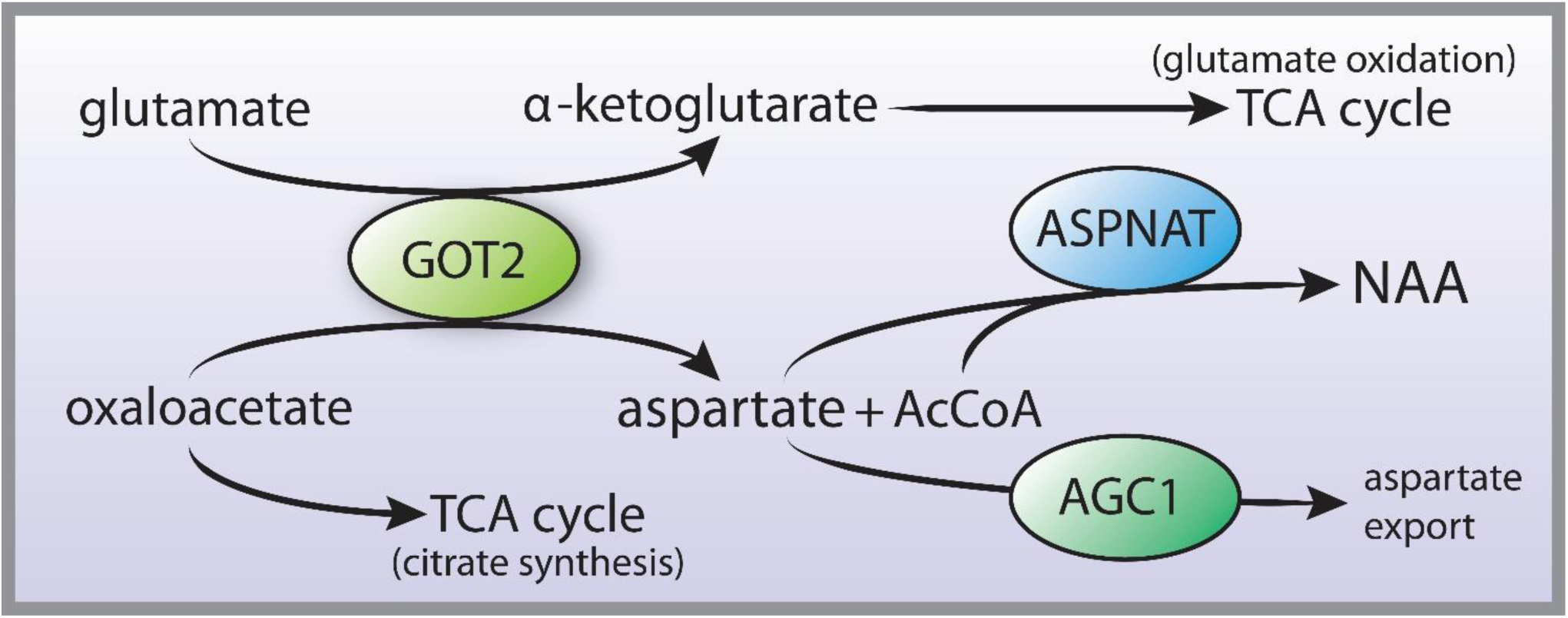
The key enzymes and transporter involved in aspartate removal from the mitochondrial matrix in neurons. The forward reaction of AAT generates *α*-ketoglutarate for oxidation in the TCA cycle and aspartate. The primary route for aspartate removal from the matrix is via the transporter AGC1 (SLC25A12 in neuronal mitochondria), which is part of the malate-aspartate shuttle. ASPNAT provides a secondary aspartate removal system whereby conversion to NAA prevents further metabolism in the matrix and allows export via an as yet undetermined dicarboxylate transporter. OAA is limiting in this reaction because the majority of OAA is consumed by citrate synthase. The remaining OAA provides substrate for the GOT2 (AAT) reaction. The model tests how this balance changes across OAA availability, export capacity, ASPNAT activity, and alternative (cytoplasmic) ASPNAT topology. Abbreviations: AGC1 = aspartate glutamate carrier 1 (aka Aralar1), ASPNAT = aspartate N-acetyltransferase, GOT2 = glutamic oxaloacetic transaminase (aka aspartate aminotransferase or AAT).

Several experimental paradigms are consistent with this framing. Isotope-exchange studies established that mitochondrial AAT readily equilibrates substrate and product pools, whereas *in vivo* and *ex vivo* ^13^C studies showed sustained routing of glutamate-derived carbon through aspartate, NAA, and related oxidative intermediates in intact brain (Cooper et al., 2023; Kimmich et al., 2002; Rodrigues et al., 2013a; Sonnewald et al., 1993). Direct synthesis measurements and tracer studies further indicate that neurons continue to make NAA despite the slow turnover of the much larger tissue reservoir, indicating continuous synthesis despite high product concentrations (Berl et al., 1970; Choi and Gruetter, 2004; Moreno et al., 2001; Truckenmiller et al., 1985). The steady rate of NAA synthesis in the face of strong product inhibition implies a thermodynamic benefit to continuous ASPNAT activity in the mitochondrial matrix.

This node is also coupled to the MAS, the main route by which cytosolic reducing equivalents are transferred into neuronal mitochondria (Ahmed et al., 2025; Borst, 2020; Brunner et al., 2026; Contreras, 2015; Palmieri and Pierri, 2010; Pardo and Contreras, 2012; Satrustegui et al., 2007b). The mitochondrial aspartate-glutamate carrier, AGC1, also known as Aralar1, exports aspartate from the matrix while returning glutamate, thereby linking aspartate disposal, redox balance, and AAT substrate supply. Because AGC1 is calcium regulated, its effective export capacity is expected to vary with neuronal activity and metabolic demand (Contreras, 2015; Hewton et al., 2021; Lindsay et al., 2014; Satrustegui et al., 2007a). Genetic loss of AGC1 strongly lowers brain aspartate and NAA levels, underscoring the coupling between mitochondrial aspartate handling and NAA metabolism (Jalil et al., 2005; Profilo et al., 2017; Ramos et al., 2011). In neurons, ASPNAT is localized to the mitochondrial matrix, but radiolabeled substrate studies have suggested that there may be an additional cytoplasmic site of synthesis (Arun et al., 2009). Mitochondrial synthesis is supported by cell-fractionation, endogenous protein, and radiolabel tracer evidence (Ariyannur et al., 2008; Ariyannur et al., 2010b; Arun et al., 2009; Pessentheiner et al., 2013; Zysk et al., 2020). For the purposes of the proposed model system, we treat mitochondrial NAA synthesis as the default route, and cytoplasmic synthesis as a possible alternate route that may be more important under certain physiological conditions. As such, we will examine both the mitochondrial and cytoplasmic routes of synthesis to determine the overall effect on the thermodynamic outcomes of the site of NAA synthesis.

Here we present a quantitative treatment of the mitochondrial aspartate node in neurons to ask four linked questions. First, can mitochondrial ASPNAT increase forward AAT throughput in a low-OAA neuronal regime? Second, is the effect graded or preferentially operative under physiological stress or high metabolic demand? Third, does the effect persist across physiological uncertainty and when OAA is allowed to vary dynamically? Fourth, how does shifting ASPNAT localization from the mitochondrial matrix to the neuronal cytoplasm affect the model system? Our aim is to provide a simplified model of the aspartate node in neuronal mitochondrial metabolism that examines some of the thermodynamic benefits of NAA synthesis.

## 2 Methods

### 2.1 System compartmentation and physiological boundary conditions

The core model was built around three dynamic mitochondrial metabolites, aspartate, *α*-ketoglutarate, and NAA. Glutamate (10 mM), OAA (160 nM), acetyl-CoA (14 *µ*M), and coenzyme A (CoA; 26 *µ*M) were treated as clamped boundary pools that represent the surrounding metabolic environment rather than dynamic state variables. The low-micromolar acetyl-CoA and CoA values are consistent with brain mitochondrial acetyl-group buffering studies (Ronowska et al., 2018; Szutowicz et al., 2013). This reduced compartmental structure isolates the mitochondrial aspartate node while retaining the metabolites most directly responsible for product pressure, substrate supply, and NAA formation, following the logic of focused compartmental and boundary-condition modeling used in prior brain-metabolism analyses (Garfinkel, 1970; Gruetter, 2009; Gruetter et al., 2001). Full parameter values, boundary concentrations, and model variants are provided in **Supplementary Tables S1, S2, S5, S8 S12, S14 and S19.**

The aspartate balance in the main model is (**equation 1**):

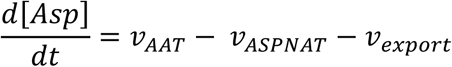

with analogous balances for *α*-ketoglutarate and NAA given in the Supplementary Information. This structure captures the competition between matrix aspartate production, ASPNAT-dependent removal, and AGC1-linked export without introducing additional whole-cell assumptions.

### 2.2 Governing kinetic rate laws

Mitochondrial AAT was represented as a reversible Haldane-consistent ping-pong bi-bi reaction, ASPNAT as an irreversible bi-substrate reaction with product inhibition by NAA and CoA, and AGC1-mediated aspartate removal as a first-order export term appropriate for the low-aspartate regime examined here (Madhavarao et al., 2003; McKenna et al., 2006a; Mutthamsetty et al., 2020). To preserve thermodynamic consistency, the reverse AAT capacity was constrained by (**equation 2**):

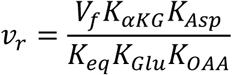

and the full kinetic expressions were evaluated in the numerical simulations reported below. Because ASPNAT activity measured in detergent-solubilized preparations underestimates native membrane-associated capacity, the effective ASPNAT parameter was treated as semi-empirical and calibrated to reproduce experimentally reported NAA synthesis rates within a biologically plausible range (Madhavarao et al., 2003; Wang et al., 2016). The complete rate laws, calibration rationale, and parameter provenance are provided in the Supplementary Methods.

### 2.3 Numerical convergence and simulation workflow

Steady states were obtained with a Livermore Solver for Ordinary Differential Equations with Automatic method switching (LSODA)-centered adaptive convergence workflow using relative tolerance 10^−10^ and absolute tolerance 10^−13^. Integration proceeded in 1000 min blocks until the maximum absolute derivative across the dynamic species satisfied max |dX/dt| < 10^−9^. For each comparison, matched ASPNAT-on and ASPNAT-off steady states were computed under the same boundary conditions. The main comparisons converged under LSODA, whereas the dynamic-OAA sensitivity extension used a stiff-solver fallback only if a simulation failed the same residual test under LSODA. Robustness analyses included Latin Hypercube Sampling across ten uncertain parameters, a correlated-sampling stress test, and a two-compartment simulated stress-rerouting test that compared the mitochondrial ASPNAT premise against an extra-mitochondrial aspartate-consuming condition while preserving the same canonical relief metric. Supplementary Methods provide the full numerical protocol, convergence criteria, parameter bounds, covariance scenarios, and extended dynamic-OAA details.

### 2.4 Relief metric definition

All reported relative effects use one canonical relief definition (**equation 3**):

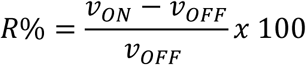

where *v* denotes net forward AAT flux for matched ASPNAT-on and ASPNAT-off simulations. Using a single denominator convention is essential because the manuscript compares baseline, robustness, and spatial-organization analyses on the same relative scale.

## 3 Results

The current model analyses were generated using parameter settings chosen a priori, not adjusted to obtain or reinforce the conclusions emphasized below. Apart from the semi-empirical ASPNAT capacity, which was constrained to remain compatible with reported NAA synthesis rates and biochemical activity bounds, the key behaviors emerged from published kinetic constraints and the assigned mitochondrial compartmentation rather than from explicit fitting of relief or regime boundaries (Madhavarao et al., 2003; McKenna et al., 2006b; Moreno et al., 2001; Truckenmiller et al., 1985; Wang et al., 2016). Transport-law substitutions, uncertainty ensembles, and structural extensions are detailed in the Supplementary Information.

### 3.1 Baseline relief identifies a product-limited mitochondrial node

In the baseline mitochondrial-compartment model, ASPNAT lowered steady-state mitochondrial aspartate from 141 to 105 µM and increased net forward AAT flux from 0.14095 to 0.184 53 nmol/min/mg, corresponding to 30.92% relief (**Figures 2A - 2C**). NAA accumulated to 7.92 mM, within the physiological range reported for neuronal NAA pools in tracer and spectroscopic studies (**Figure 2C**), while the modeled state remained close to thermodynamic reversal (Γ/Keq = 0.9206) (Müller et al., 1994; Rodrigues et al., 2013b). The baseline comparison points to a clear mechanism; ASPNAT does not open a new high-flux route, it trims the matrix aspartate pool enough to move AAT away from a product-limited state.

**Figure 2.**
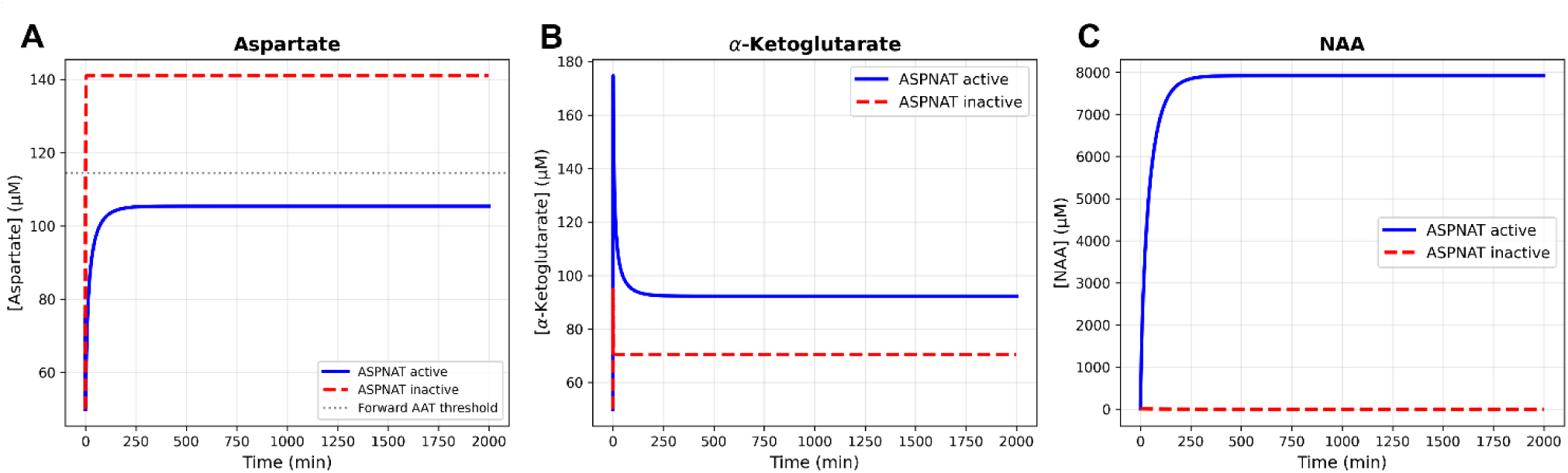
Baseline ASPNAT activity relieves the product-limited mitochondrial aspartate node. This comparison defines the reference low-OAA operating point used in the current model comparing ASPNAT-on versus ASPNAT-off states. A-C) In the paired simulations, ASPNAT lowers matrix aspartate, increases *α*-ketoglutarate concentration, and builds an NAA pool accompanying the 30.9% increase in forward AAT flux.

The baseline trajectories show a coordinated state shift rather than a single isolated flux change. When ASPNAT is turned on, matrix aspartate falls, α-ketoglutarate shifts modestly upward, and NAA accumulates as the new sink fills. That pattern is what one expects from a product-limited AAT node. ASPNAT does not create a bypass reaction; it changes the balance of the existing reaction pair by removing one of the products that keeps the mitochondrial state close to reversal.

### 3.2 Local sensitivity points to thermodynamic control

The local sensitivity analysis supports the same mechanistic conclusion from an independent direction (**Figures 3A, B**). For AAT flux, the strongest positive controllers were OAA, glutamate, the AAT equilibrium term, and downstream α-ketoglutarate removal, each with scaled sensitivities of about +0.4 in the baseline state. AGC1-mediated export also exerted substantial positive control on forward AAT flux. By contrast, AAT catalytic capacity itself had near-zero local control (S ≈ +0.04). This pattern says the node is constrained mainly by thermodynamic headroom and product disposal, not by a shortage of AAT enzyme activity.

**Figure 3.**
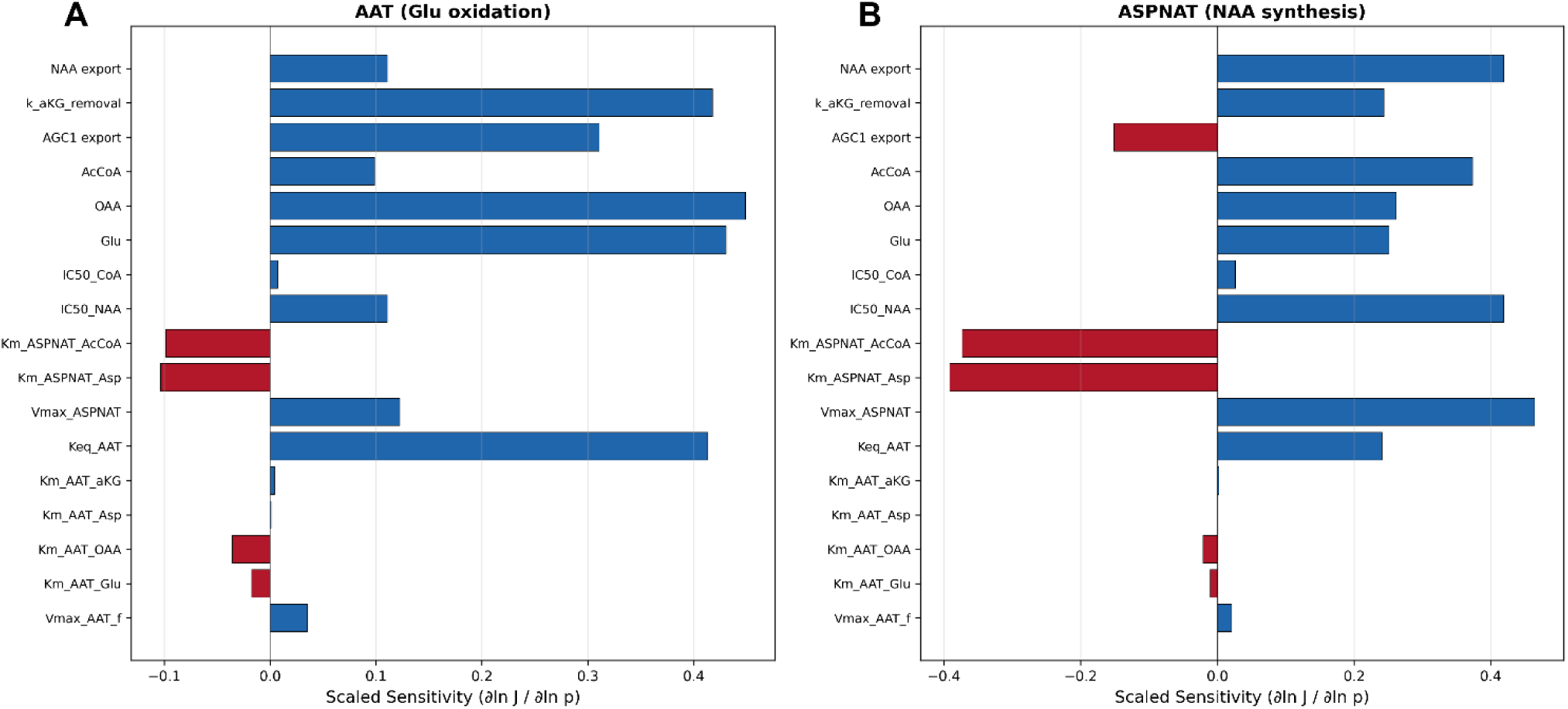
Local sensitivity identifies thermodynamic control parameters. A) The scaled local sensitivities for forward AAT throughput show primary control by OAA availability, the AAT equilibrium term, downstream *α*-ketoglutarate removal, and AGC1-linked export. B) The scaled local sensitivities for ASPNAT flux show primary control by ASPNAT capacity, product inhibition, and acetyl-CoA supply.

Assuming mitochondrial ASPNAT localization, the baseline mitochondrial aspartate node sits close to AAT reversal, which is why small shifts in matrix aspartate have outsized consequences for forward flux (**Figures 4A, B**). This spatial premise shapes the relief mechanism and is tested explicitly below. Within that simulated condition, the reference operating point lies adjacent to the reversal boundary rather than deep inside a comfortably forward regime. A low-capacity irreversible sink can therefore exert disproportionate leverage because product removal moves a reaction pair that is already poised near equilibrium.

**Figure 4.**
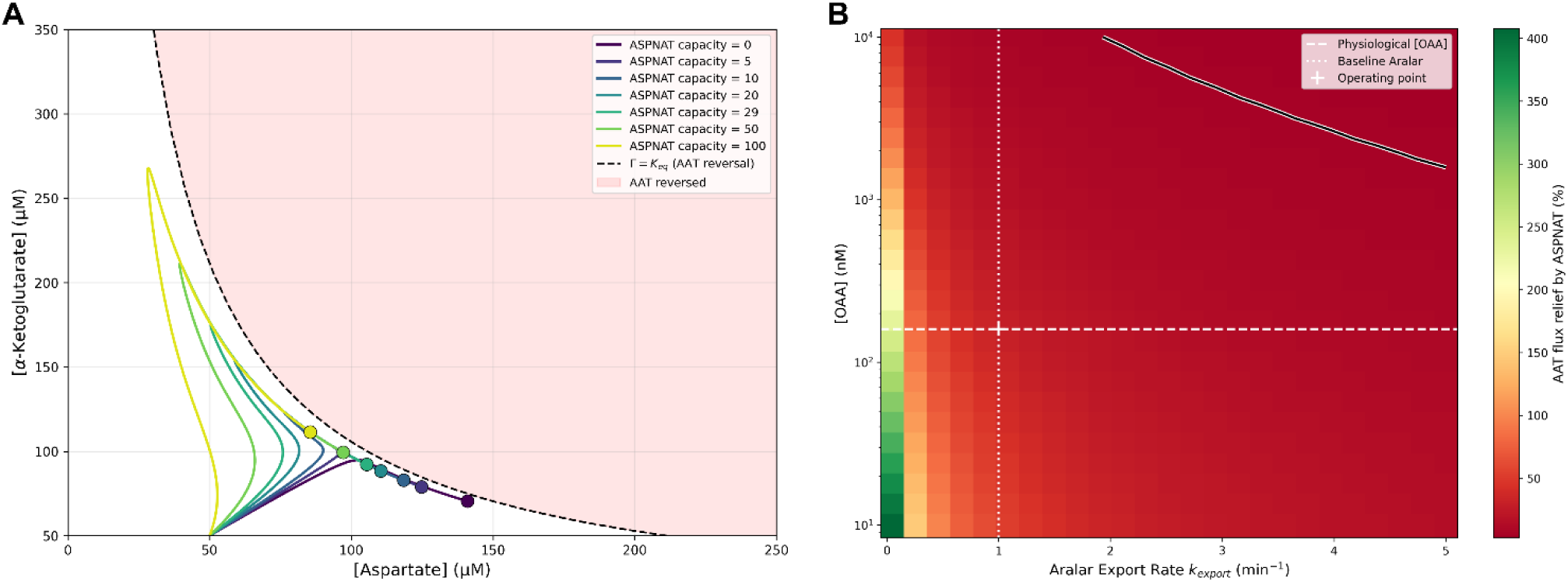
The mitochondrial aspartate node sits close to AAT reversal, creating a vulnerability that mitochondrial ASPNAT preferentially relieves in a constrained low-OAA, weak-export regime. These panels assume mitochondrial-localization. A) The phase-plane view places the reference operating point close to the AAT reversal boundary (dashed line). B) The OAA x AGC1 heat map shows that relative AAT relief is largest in the low-OAA, low-export regime and drops to 7% once OAA becomes abundant or export is restored. ASPNAT provides a greater thermodynamic assist when key substrates are limited.

This baseline geometry explains how the relief mechanism works, but not when it matters physiologically. Whether ASPNAT contributes meaningfully depends on the surrounding state of the node, especially OAA supply and the strength of the primary AGC1-linked export route that clears aspartate from the matrix.

### 3.3 Dynamic OAA preserves the directional relief effect

The fixed-OAA baseline isolates the thermodynamic mechanism most cleanly, but OAA is not a static metabolite in living cells. We therefore relaxed the OAA boundary in a dynamic sensitivity extension with malate dehydrogenase and citrate synthase terms (**Supplementary Figure S1**). The accepted simulation preserved the direction of ASPNAT-mediated AAT relief, with 33.98% relative relief under the same canonical denominator. The result should not be construed as a physiological flux prediction, because the operating flux of the reduced extension is subphysiological and the ASPNAT-off simulation required stiff-solver verification. The sign of the relief effect did not disappear when the fixed-OAA clamp was relaxed.

### 3.4 Low OAA, acute AGC1 restriction, and ASPNAT capacity define a graded relief regime

The bivariate OAA × acetyl-CoA analysis distinguishes substrate-supported NAA synthesis from the thermodynamic advantage (**Figures 5A – 5D**). Absolute ASPNAT flux increased with acetyl-CoA and OAA availability, indicating that NAA synthesis is driven by substrate supply. The residual aspartate export fell as ASPNAT substrate supply increased, which corresponds with experimental data on NAA efflux from isolated brain mitochondria (Patel and Clark, 1979). By contrast, relative AAT relief scaled with decreasing OAA and increasing acetyl-CoA concentrations. The dependence of the AAT relief effect on acetyl-CoA availability can be seen in **Figure 5D**, where low oxaloacetate has little effect on the relief at low acetyl-CoA levels, but strong effects at high acetyl-CoA concentrations.

**Figure 5.**
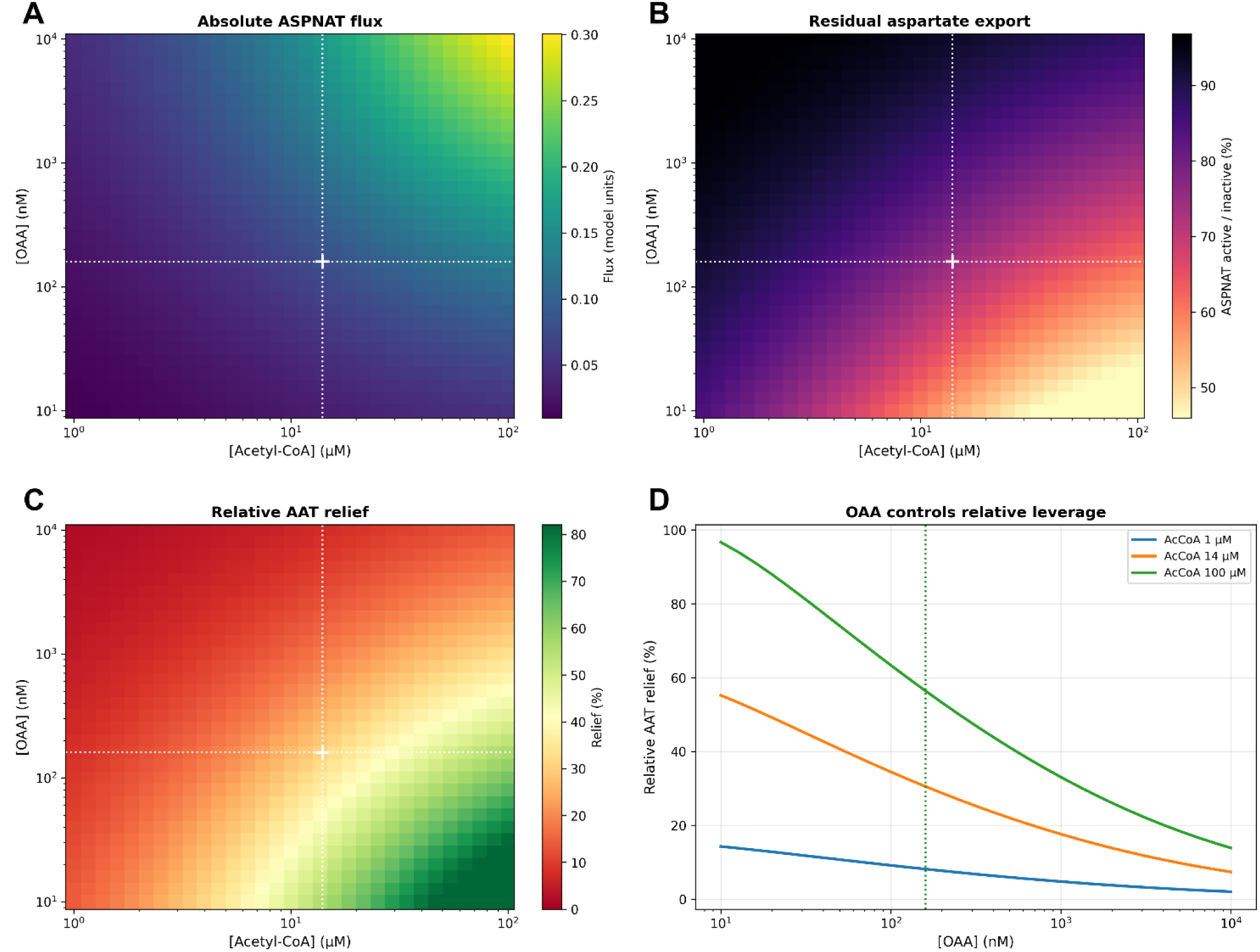
Heat maps and graph showing how oxaloacetate and acetyl-CoA concentrations affect ASPNAT flux, aspartate export and AAT relief. These OAA *×* acetyl-CoA sweeps use clamped boundary pools to isolate local node behavior; they do not model dynamic acetyl-CoA depletion. A) Absolute ASPNAT flux increases with increased OAA and acetyl-CoA availability. B) Residual aspartate export falls as OAA levels drop and acetyl-CoA levels increase. C) Relative AAT relief is strongest in the low-OAA regime and increases steadily as acetyl-CoA concentration is increased. D) The OAA concentration steps show that acetyl-CoA supports the AAT relief effect, whereas low OAA defines the thermodynamic state in which that sink has the largest proportional effect. Dashed lines indicate physiological values.

To examine how oxaloacetate concentrations and the AGC1 export rate for aspartate affect system parameters, we used one-dimensional sweeps to examine system component responses (**Figure 6**). As expected, AAT flux and aspartate levels increase with increasing OAA concentrations (**Figures 6A, B**). When OAA is scarce, the relative ASPNAT enhancement effect is large because the node is thermodynamically sensitive to product removal. As OAA rises, absolute forward AAT flux becomes less product-limited and the incremental ASPNAT relief contracts toward minimal values. At the estimated low-OAA reference condition of 160 nM, the model remains well inside the relief-favored regime, with the effect remaining around 31% (**Figure 6C**). This proportional leverage is distinct from absolute ASPNAT or NAA synthetic capacity, which depends on acetyl-CoA supply and is evaluated separately in **Figure 5** and Supplementary **Table S20**.

**Figure 6.**
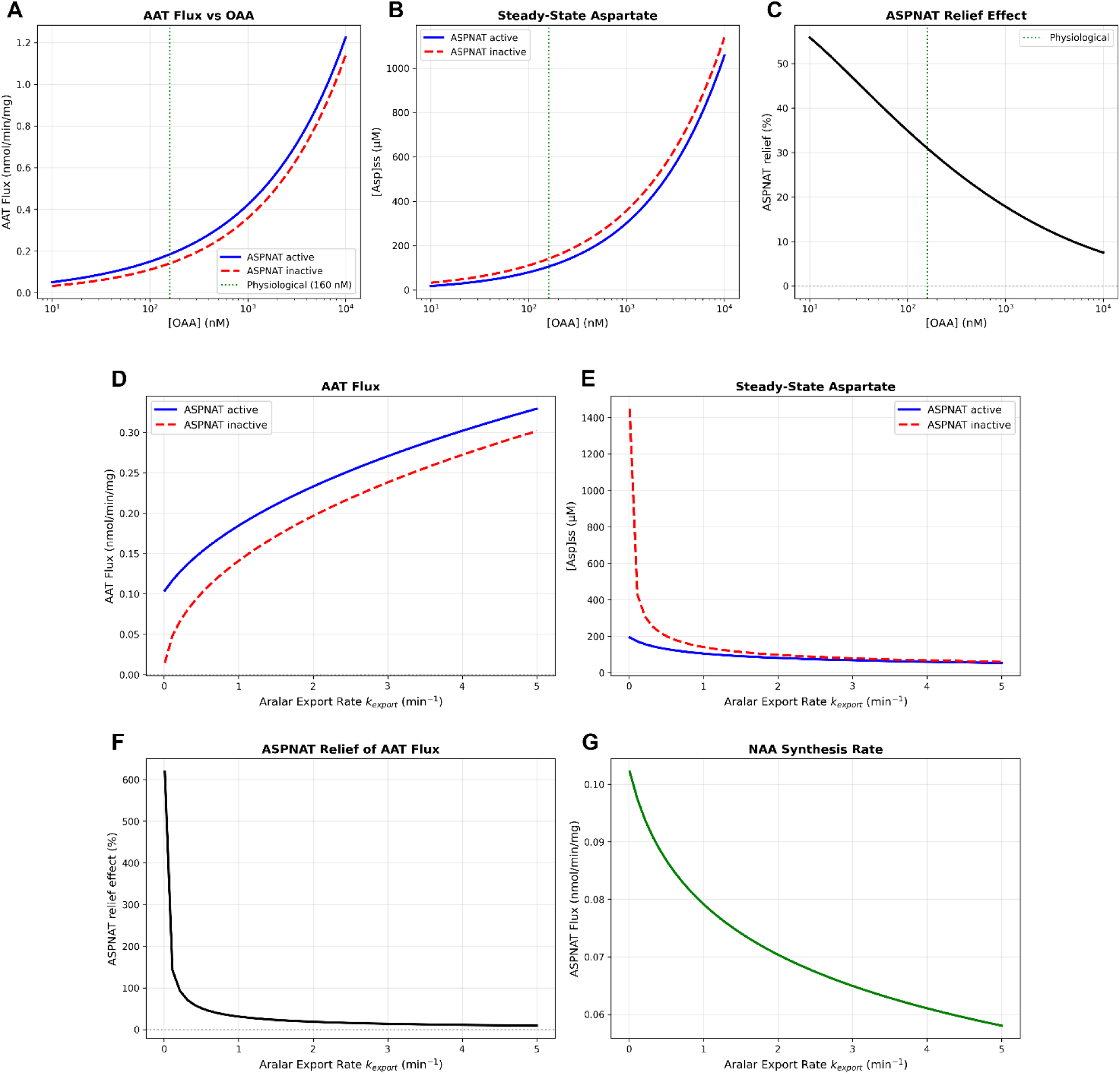
ASPNAT provides auxiliary relief when OAA supply is limited or AGC1 export is acutely restricted within the constrained operating regime. These one-parameter sweeps probe local node dynamics around the estimated low-OAA reference condition rather than whole-neuron fluxes. A-C) The OAA sweeps report forward AAT flux, steady-state matrix aspartate, and the relative ASPNAT relief metric (dashed green lines show OAA physiological concentration). D-G) The AGC1 (Aralar) export sweeps report forward AAT flux, steady-state matrix aspartate, relative ASPNAT relief, and NAA synthesis rate.

The effect of aspartate export by AGC1 (Aralar1) on AAT transamination flux showed a positive correlation and ASPNAT activity shifted the AAT flux upward (**Fig 6D**). ASPNAT activity is critical for keeping matrix aspartate levels in check when AGC1 activity is low (**Fig 6E**). When aspartate export is weak, ASPNAT provides an auxiliary removal path and the relief effect rises. When export is restored, the primary route clears matrix aspartate efficiently and the ASPNAT contribution is greatly reduced (**Fig 6F**). These sweeps therefore place ASPNAT in an auxiliary role rather than treating it as a dominant controller of the node. As AGC1 aspartate export increased, NAA synthesis by ASPNAT is decreased (**Fig 6G**).

Within that constrained regime, the ASPNAT response remains graded and concentration-dependent (**Figures 7A-D**). Increasing ASPNAT activity from 0 to 300% of the baseline value raised forward AAT flux monotonically from 0.141 to about 0.22 nmol/min/mg, lowered steady-state mitochondrial aspartate from 141 µM to 88 µM, and expanded the modeled NAA pool toward approximately 13 mM. These results show that ASPNAT has substantial effects on this node in mitochondrial metabolism, facilitating aspartate removal from the matrix, maintaining AAT forward flux and providing assistance to AGC1.

**Figure 7.**
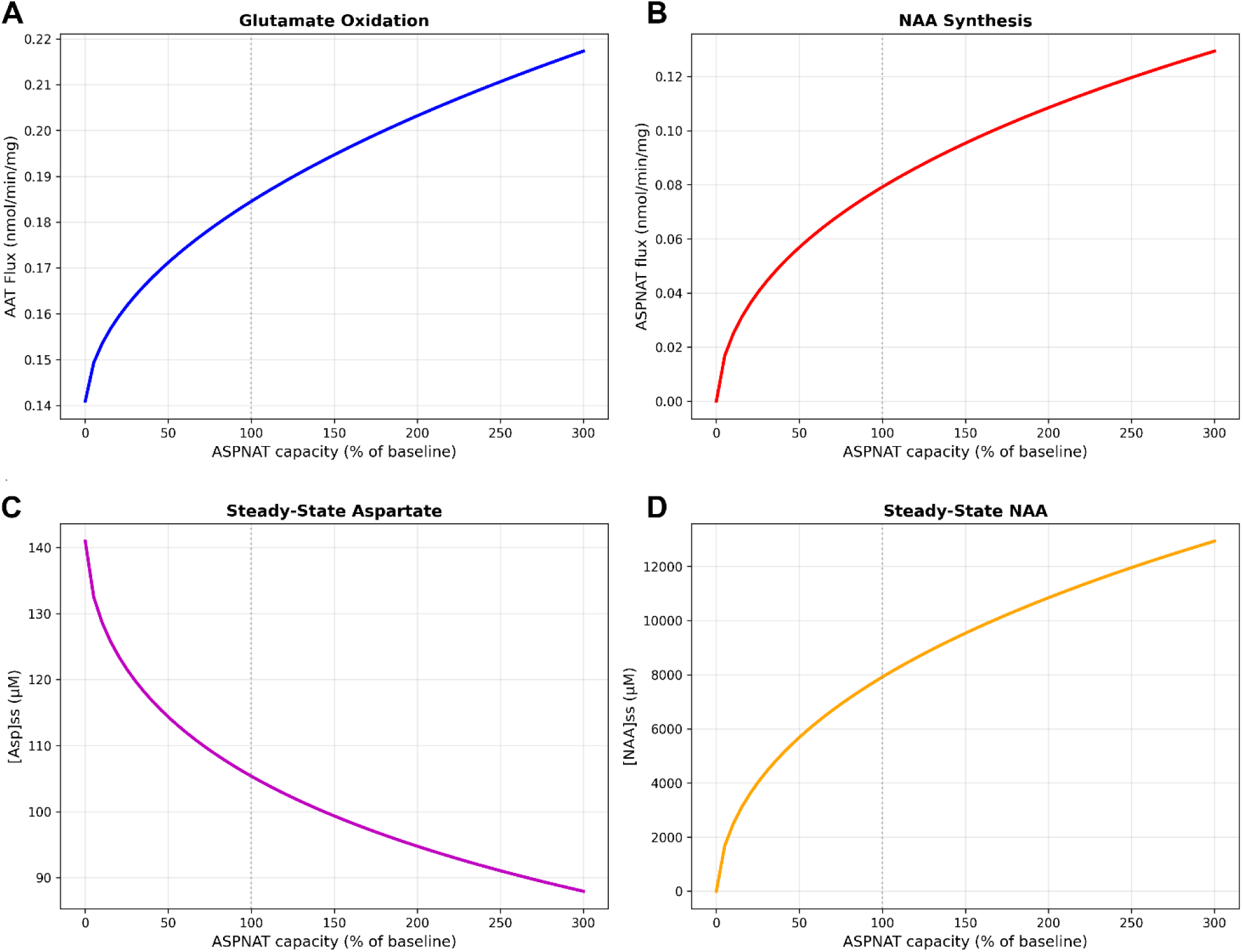
ASPNAT capacity titration produces graded changes rather than all-or-none responses. A) Forward AAT flux rises progressively as ASPNAT capacity increases and B) ASPNAT flux scales with available enzyme capacity. C) Mitochondrial aspartate falls continuously across the titration and D) the modeled NAA pool expands as the ASPNAT aspartate sink strengthens.

### 3.5 ASPNAT-mediated aspartate removal requires acetyl-CoA and incurs a modest local carbon cost

The capacity of ASPNAT to facilitate AAT throughput is still limited by substrate availability because ASPNAT cannot operate without a continuous supply of acetyl-CoA (**Figures 5 and 7**). When the clamped mitochondrial acetyl-CoA boundary pool was titrated from 1 µM to 100 µM, NAA synthesis and relative relief both increased smoothly (**Figure 5D**). At the estimated physiological value of about 14 µM for acetyl-CoA, the pathway was active but not saturated. At lower acetyl-CoA, ASPNAT velocity fell sharply and the relief effect contracted because enzyme activity became highly substrate-limited.

We next compared matched ASPNAT-on and ASPNAT-off steady states to determine the energetic cost of operating the AAT relief valve (**Figures 8A-F**). The comparison is deliberately local. It reports comparative proxy indices linked to forward AAT throughput, MAS support, cytosolic NAD+ regeneration, ATP-linked throughput, ATP fraction, acetyl-CoA drain, and the effective acetyl-CoA to CoA ratio. It does not estimate a whole-neuron ATP or redox budget.

**Figure 8.**
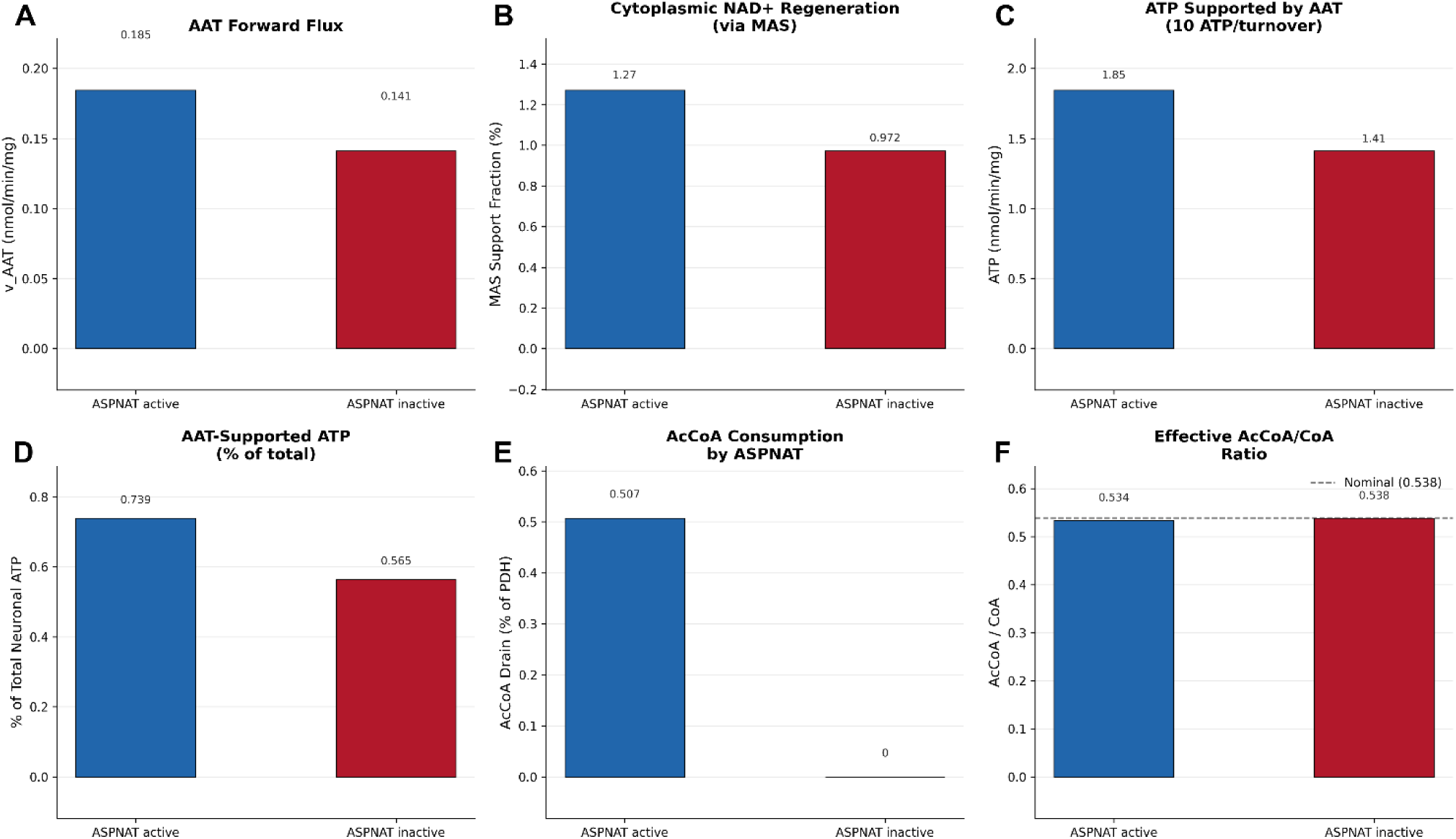
Local energetic proxy comparison for the ASPNAT-active and ASPNAT-inactive baseline states. The bar plots use matched steady states and are intended as comparative indices, not as a whole-neuron ATP or redox budget. A) Forward AAT flux is preserved by the ASPNAT sink (0.185 vs 0.141 nmol/min/mg). B) The MAS-linked cytosolic NAD+ regeneration proxy shifts in the same direction (1.27 vs 0.972%). C) AAT-linked ATP throughput increases under the local proxy calculation (1.85 vs 1.41 nmol/min/mg). D) The corresponding ATP fraction remains small on the whole-neuron scale (0.739 vs 0.565%). E) ASPNAT imposes a modest acetyl-CoA drain relative to the represented oxidative context (0.507% of PDH). F) The effective acetyl-CoA to CoA ratio remains close to the nominal boundary-pool ratio (0.534 vs 0.538).

Within that comparative frame, the ASPNAT-on state enhanced forward AAT flux (**Figure 8A**) and shifted redox-linked indices in the same favorable direction (Figs 8B-D), while imposing only a modest fractional acetyl-CoA drain relative to baseline (**Figures 8E, F**). Taken together, the panels in **Figure 8** suggest a synergistic linkage between ASPNAT and AAT. ASPNAT gains leverage over a near-equilibrium AAT step by spending acetyl-CoA to trap aspartate irreversibly. The mechanism is useful because ASPNAT-mediated product removal preserves near-equilibrium forward AAT flux, and it is vulnerable because that safeguard depends on acetyl-CoA availability. The current clamped-boundary model does not test dynamic acetyl-CoA depletion.

### 3.6 ASPNAT extra-mitochondrial localization reverses the sign of the predicted metabolic effect

Our model assumes ASPNAT localization to the inner surface of the mitochondrial inner membrane, facing the matrix. (**Figure 9**). However, radiolabel incorporation experiments with an inhibitor of AAT, aminooxyacetic acid, suggest that there may be a cytoplasmic source of NAA synthesis (Arun et al., 2009), which could be more operative under physiological stress. Under the mitochondrial premise, net forward AAT flux increased by 53.32%. In the extra-mitochondrial aspartate-consuming simulation condition, the sign reversed and forward AAT flux fell by 17.80% (**Figure 9A**). The reason is that mitochondrial ASPNAT removes aspartate directly from the matrix, whereas an extra-mitochondrial aspartate sink does little to affect matrix levels. Mitochondrial localization provides multiple, often subtle metabolic benefits that do not accrue when the localization is situated extra-mitochondrially. Mitochondrial ASPNAT lowers matrix aspartate (**Figure 9B**) and increases cytoplasmic aspartate when ASPA is present (**Figure 9C**). Extra-mitochondrial ASPNAT localization negates the AAT relief effect and reduces cytoplasmic aspartate levels (**Figure 9D**). It is currently unknown the extent to which cytoplasmic NAA synthesis occurs under normal physiological conditions, or if pathology might shift NAA synthesis away from the matrix. Evidence from hepatocellular carcinoma cells indicates that reducing ASPNAT activity increases cytoplasmic aspartate levels, indicating that the site of ASPNAT activity is mitochondrial, rather than cytoplasmic (De Falco et al., 2023). Further studies are required to determine if certain pathologies present conditions that shift NAA synthesis from the mitochondrial matrix to the cytoplasm.

**Figure 9.**
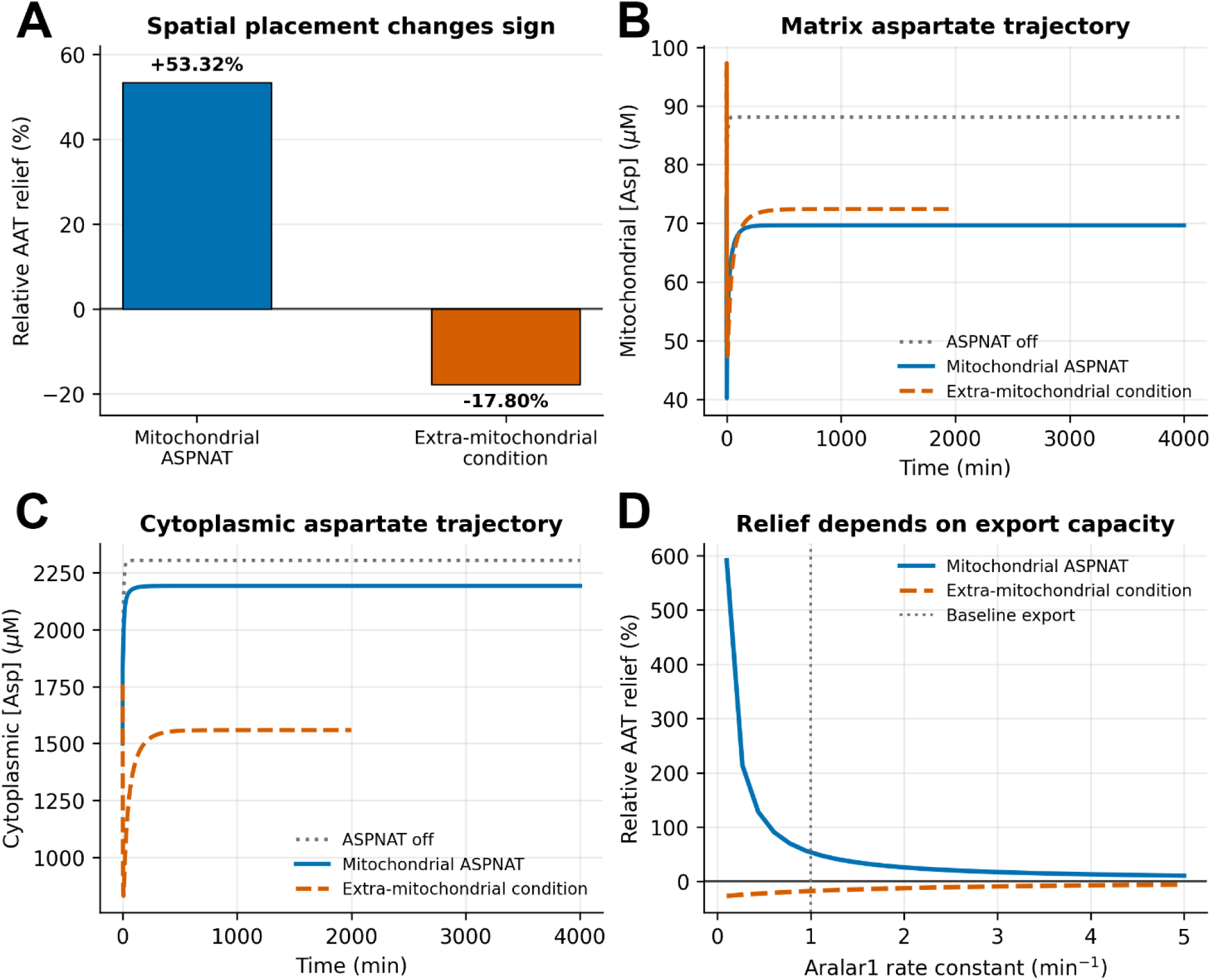
Simulated extra-mitochondrial aspartate consumption reverses the sign of the metabolic response. Mitochondrial ASPNAT is the supported biological premise; the extra-mitochondrial condition is included only as a falsifiable simulation of pathological or stress-related rerouting of effective aspartate consumption. A) Mitochondrial ASPNAT increases forward AAT flux by 53.32%, whereas the extra-mitochondrial condition reverses the sign of the relief metric and decreases forward AAT flux by 17.80%. B) The time course shows mitochondrial aspartate concentration during the spatial-placement simulations. C) The time course shows cytoplasmic aspartate concentration during the same simulations. D) The AGC1 (Aralar1) rate-constant sweep shows how the relief effect changes as primary aspartate export capacity varies.

### 3.7 Gene deficiency effects on NAA synthesis and AAT relief effect

We investigated the effects of gene deficiencies on NAA synthesis, the AAT relief effect and AAT flux. The model employed the three species of the mitochondrial aspartate node: aspartate, α-ketoglutarate, and NAA, with fixed pools of glutamate, OAA, acetyl-CoA, and coenzyme A (**Figure 10**). The conditions evaluated included complete loss of ASPNAT activity, a 50% reduction in AGC1 activity and a 60% reduction in ASPA activity. The reduced ASPA condition uses residual NAA clearance calibrated to produce a heightened NAA pool in the range observed in Canavan disease.

**Figure 10.**
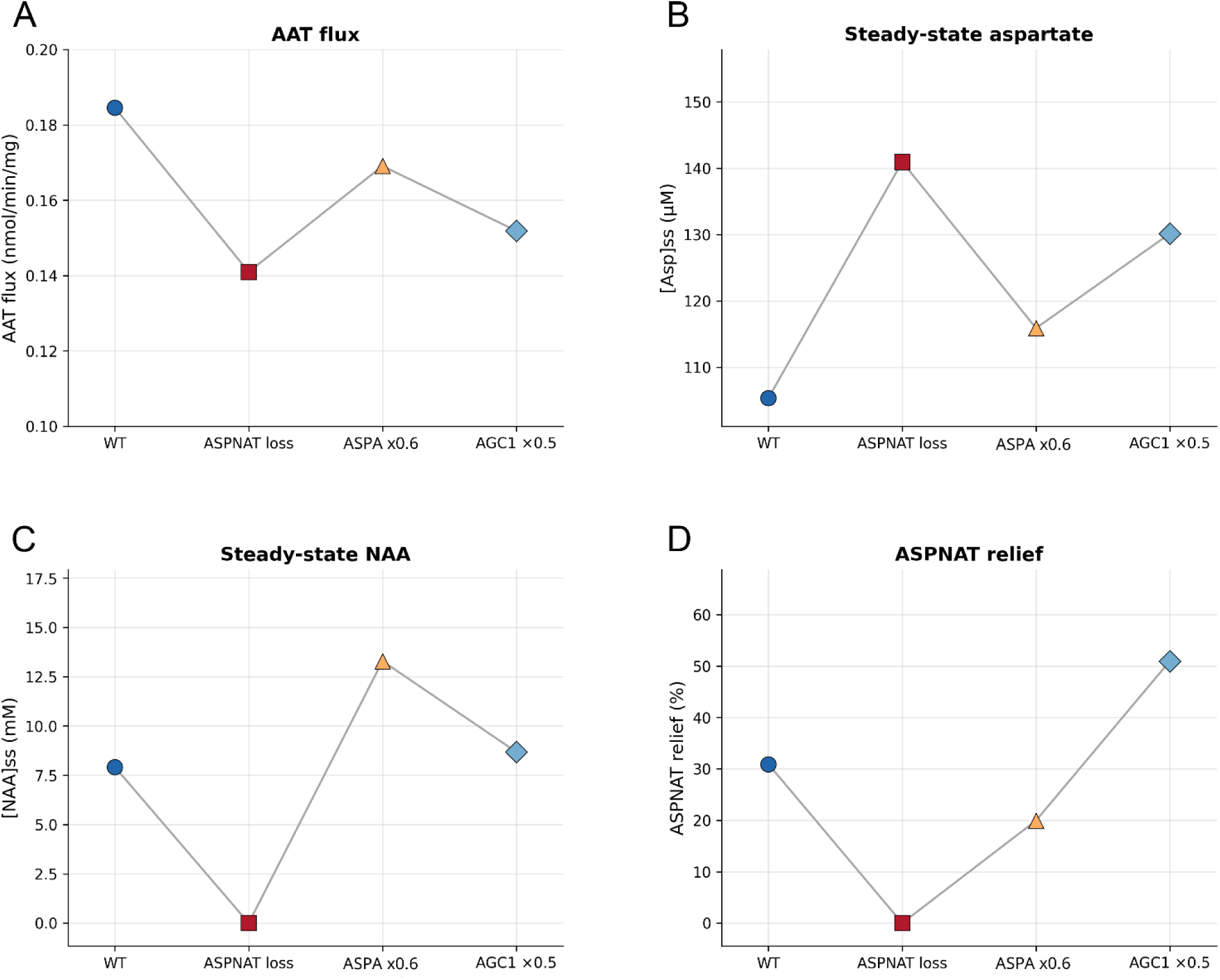
Steady-state AAT flux, aspartate and NAA pools, and relief (R%) for wild-type mitochondria and three deficiency conditions: ASPNAT loss, 60% ASPA deficiency, and a 50% reduction in AGC1-mediated aspartate export.

AAT flux fell markedly under ASPNAT deficiency: with ASPNAT active, flux was 0.185 nmol · min⁻¹ · mg⁻¹, whereas turning ASPNAT off reduced it to 0.141 nmol · min⁻¹ · mg⁻¹, corresponding to a 30.9% decrease (**Fig 10A**). Without ASPNAT, relief falls to zero (**Fig 10D**) as this removes the sink for matrix aspartate. In the low-OAA neuronal regime (160 nM), forward AAT is product-limited; without acetylation, aspartate accumulates, the reaction drifts toward equilibrium, and relief disappears.

ASPA activity reduction to 50% of wild type reduced AAT flux (**Fig 10A**) and increased intramitochondrial aspartate (**Fig 10B**). Under the simulated partial ASPA deficiency (NAA clearance reduced until the steady pool approaches the published Canavan level of 13.7 mM), NAA reaches 13.3 mM and R% falls to 20 % as product inhibition restrains ASPNAT (**Fig 10D**). When ASPA cannot hydrolyze NAA, it accumulates and inhibits ASPNAT (IC_50_ 0.85 mM), so synthesis slows.

When AGC1 export is halved, R% rises to 51 % because matrix aspartate builds up and drives AAT forward by mass action (**Fig 10D**). This restricts aspartate export from the matrix and the trapped aspartate then pushes AAT forward and inflates R%. That pattern marks a limitation of the three-pool model. A two-compartment model with cytosolic GOT1 is needed to capture the full AGC1 phenotype.

## 4. Discussion

### 4.1 NAA synthesis depends on anaplerotic substrate availability

Our model indicates that NAA synthesis in neuronal mitochondria has positive thermodynamic effects under certain conditions. The AAT relief effect increases when the aspartate node is simultaneously close to AAT reversal, poor in OAA, and insufficiently cleared by the primary AGC1 export route. Under those combined constraints, removing matrix aspartate changes the thermodynamic margin more effectively than increasing AAT catalytic capacity itself. Outside that regime, the aspartate relief effect wanes and NAA synthesis shifts to a parallel pathway to citrate for exporting an acetyl-CoA precursor from the mitochondrial matrix to the cytoplasm in order to support lipid synthesis.

During high substrate availability, especially when aspartate and acetyl-CoA are sufficient, absolute NAA synthesis can be high and the acetate moiety supports downstream lipid-related carbon incorporation. Under those high-substrate conditions, the relative aspartate relief effect is lower because OAA deficiency and AAT product limitation are less severe. The aspartate relief effect is most apparent when low OAA places AAT near reversal, even though absolute NAA synthesis can be substrate-limited in that regime. The relief effect can remain during high throughput, but it is greatly reduced in relative magnitude.

This conditional architecture fits earlier brain metabolism work in which aspartate handling, MAS function, and glutamate oxidation are tightly coupled modules (Borst, 2020; Pardo and Contreras, 2012; Satrustegui et al., 2007a; Yudkoff et al., 1994). MAS shuttle activity is tied directly to aspartate availability in the mitochondrial matrix (Brunner et al., 2026). NAA synthesis is also dependent on aspartate levels, which in turn is dependent on oxaloacetate levels via the action of AAT. Because oxaloacetate levels in the mitochondrial matrix are in the nanomolar range, the aspartate node modeled here operates within a range where the NAA relief effect on AAT throughput is meaningful.

### 4.2 AGC1 competes with ASPNAT and Citrate Synthase for substrate

The mitochondrial matrix concentration of OAA is a limiting factor in mitochondrial throughput. The low oxaloacetate concentration impacts on matrix aspartate concentration by limiting AAT flux. The aspartate generated by AAT has 3 main clearance routes; export via AGC1, or conversion to either NAA by ASPNAT, or citrate by the action of citrate synthase. These 3 systems compete for a scarce resource. In the model presented here, lowering AGC1 export capacity asks what happens when a primary matrix-aspartate clearance route is transiently weak, as might occur in a low-calcium or low-shuttle-demand state. The AGC1-mediated aspartate export sweep therefore answers the question of when an auxiliary sink becomes useful. NAA synthesis is maximal when substrate levels are high, but even at lower substrate supply levels, NAA synthesis facilitates AAT throughput by lowering product inhibition.

### 4.3 NAA synthesis provides local enzymatic relief and acetyl-group transfer

Whole-brain NAA turnover appears slow on the scale of hours to days (Choi and Gruetter, 2004; Xu et al., 2008)), but the mechanism modeled here acts at the mitochondrial site of synthesis, where instantaneous removal of matrix aspartate can change the thermodynamic margin of a near-equilibrium AAT reaction. The ASPNAT synthetic step drains a strategically positioned substrate pool quickly enough to influence the node.

The seminal acetate-transfer and mitochondrial efflux studies of D’Adamo, Patel-Clark and others showed that the acetyl group in NAA can participate in lipid-related carbon handling (D’Adamo and D’Adamo, 1968; D’Adamo and Yatsu, 1966; Madhavarao et al., 2005; Mehta and Namboodiri, 1995; Moreno et al., 2001; Patel and Clark, 1979, 1980, 1981). Our model investigates how NAA synthesis facilitates AAT throughput, especially when AAT is nearing reversal conditions. These distinct actions, acetyl group transfer and thermodynamic assist, are closely tied together at a central node in mitochondrial metabolism where the fate of the acetate moiety of acetyl-CoA is decided by either citrate lyase or ASPNAT. The addition of ASPNAT as a complementary pathway to this key step in carbon handling in the citric acid cycle provides additional control and throughput.

### 4.4 The relief mechanism is coupled to acetyl-CoA availability

When mitochondrial matrix acetyl-CoA is adequate, ASPNAT can trap aspartate irreversibly and preserve forward AAT handling. When acetyl-CoA falls, enzyme activity becomes substrate-limited and the relief effect weakens. Literature on brain acetyl-group buffering already suggests that low-micromolar mitochondrial acetyl-CoA pools and acetylcarnitine-carnitine buffering can redistribute acetyl units under stress (Ferreira and McKenna, 2017; Ronowska et al., 2018; Szutowicz et al., 2013). In conditions such as hypoxia, ketosis, or other redox-shifted states, oxaloacetate availability and acetyl-CoA availability may change together, which would narrow the operating window for ASPNAT-dependent relief (Clanton et al., 2017). The mitochondrial matrix has the highest acetyl-CoA level of any cellular compartment (Wang et al., 2023) and our model predicts that ASPNAT-mediated enhancement of AAT throughput is coupled to a measurable, but very limited, local acetyl-CoA cost. During times of high metabolic activity, the lactate shuttle between astrocytes and neurons directs lactate generated in astrocytes to neurons, which convert the lactate back to pyruvate. Neurons can then convert the pyruvate to acetyl-CoA to enter the TCA cycle, providing additional acetyl-CoA for oxidation in neuronal mitochondria. This could be one reason why neurons benefit from having another output pathway for the high acetyl-CoA levels in the mitochondrial matrix.

### 4.5 NAA declines due to injury are consistent with metabolic reprioritization

Multiple lines of evidence indicate that injury-induced reductions in NAA concentrations in the brain are evidence of neuronal injury or loss (reviewed in Moffett et al., 2013). Early NAA declines following brain injury are also consistent with a reversible metabolic reprioritization in which falling acetyl-CoA availability reduces NAA synthesis and starves the relief valve before overt cellular loss is apparent.

NAA levels fall rapidly in disorders that impose metabolic stress on neurons, including hypoxic–ischemic insults, mitochondrial defects, and neurodegenerative disease states (Kori et al., 2016; Moffett et al., 2013; Sharma et al., 2010; Urquiza et al., 2026; Wisnowski et al., 2016). The model demonstrates that declines in key mitochondrial metabolite levels will impact on both mitochondrial throughput and NAA synthesis. The NAA decline can result from structural injury, decreased synthesis and accelerated NAA hydrolysis. For example, post-injury aspartoacylase (ASPA) upregulation could contribute to NAA decline and may redirect acetate toward repair-associated lipid synthesis rather than reflecting ASPNAT limitation alone (Moffett et al., 2013; Moffett et al., 2020a). What the model provides is a concrete mechanism by which declining NAA can track metabolic fragility as well as neuronal integrity. If these mechanisms are correct, then an early fall in NAA can indicate a substrate-constrained bioenergetic state, increased catabolism, and mitochondrial dysfunction, rather than serving only as a marker of irreversible neuronal loss.

Early reports demonstrated that ASPA was localized primarily to oligodendrocytes in the CNS (Kirmani et al., 2002; Klugmann et al., 2003). Subsequently, using two different, well-characterized antibodies, the expression of ASPA was extended to microglia throughout the brain, as well as to neuronal groups and axonal fibers in the brainstem and spinal cord (Madhavarao et al., 2004; Moffett et al., 2011). The localization of ASPA in specific neuronal groups and their axons indicates that in those neurons, NAA can be both synthesized and catabolized in a single cell type in the CNS. In this situation, modeling NAA synthesis and breakdown is greatly simplified.

### 4.6 The predicted sign reversal resulting from cytoplasmic ASPNAT localization provides a falsifiable test for metabolic rerouting

Evidence for the localization of ASPNAT to the mitochondrial matrix is convincing. Endogenous adipocyte fractions, and neuronal mitochondrial assays support mitochondrial ASPNAT (Ariyannur et al., 2010a; Arun et al., 2009; Patel and Clark, 1979; Pessentheiner et al., 2013; Zysk et al., 2020). Loss of ASPNAT function would reduce forward AAT-linked throughput and worsen the local redox-linked activity of the MAS. If injury or disease states shift effective ASPNAT-linked aspartate consumption to the cytoplasm, our model suggests that it would entail a loss of the thermodynamic benefit of mitochondrial localization. The model indicates that an increased metabolic burden may accompany shifting NAA synthesis to the cytoplasm, but increased demand for lipid synthesis after injury may outweigh the imposed burden on mitochondrial throughput. Radiolabel or mass-label metabolite tracing studies in intact and injured brain could distinguish whether NAA synthesis shifts from a mitochondrial to a cytoplasmic site. The use of potent bisubstrate ASPNAT inhibitors and related chemical probes are available to assist these studies (Mutthamsetty et al., 2020). The model proposes a mitochondrial AAT-relief hypothesis with a falsifiable pathology-related rerouting of NAA synthesis.

### 4.7 Adipose tissue is a plausible extension of the model

Adipose tissue, especially brown adipose tissue (BAT), expresses ASPNAT, which has been localized to the mitochondria in these cells (Pessentheiner et al., 2013). Further, using ASPNAT-silenced adipocytes and BAT tissue from ASPNAT-KO mice, these investigators also found that the expression of ATP-citrate lyase is increased, indicating a compensatory mechanism to sustain acetyl-CoA throughput when the NAA route is blocked. These findings support the idea that ASPNAT participates in acetyl-group transfer and lipid synthesis-related metabolism in a single cell type, not only in the neuronal-oligodendrocyte division of labor emphasized in brain.

The extension to adipocytes must remain qualitative at this stage. Adipocytes differ from neurons in substrate supply, acetyl-CoA handling, redox demand, thermogenic state, and mitochondrial transport. For example, uncoupling proteins UPC2 and UCP3 in adipocytes can transport aspartate from the mitochondrial matrix (De Leonardis et al., 2024; Vozza et al., 2014), adding additional complexity not modeled here. UCP2 is also expressed in the CNS, especially in glial cells, but does not transport aspartate out of brain mitochondria (McKenna et al., 2006b). Therefore, the present parameter values could not be applied directly to adipocytes. Nonetheless, the ASPNAT sink could, in principle, also relieve a product-limited AAT/MAS node in adipose tissue, but modeling this would require adipocyte-specific constraints be applied.

### 4.8 Does NAA just reflect mitochondrial energy status, or does it contribute to it as well?

The earliest studies on NAA synthesis showed that NAA was only synthesized in brain mitochondria when glucose was provided (Buniatian et al., 1965) and when aspartate and acetyl-CoA were available (Patel and Clark, 1979). Based on these observations, the question arose: does NAA just reflect energy status, or does it contribute to it as well? Numerous lines of evidence link NAA synthesis and catabolism with mitochondrial energetics and energy metabolism in general. For example, in isolated brain mitochondria, inhibitors of all 4 mitochondrial transport chain complexes reduce NAA synthesis, oxygen consumption and ATP synthesis in parallel (Bates et al., 1996). Mild to moderate traumatic brain injury reduces ATP and NAA levels reversibly, and the loss and recovery of both metabolites follows a similar time course (Di Pietro et al., 2014; Signoretti et al., 2008; Vagnozzi et al., 2007). Overexpression of ASPNAT in immortalized brown adipogenic cells leads to increased mitochondrial mass, number, and oxygen consumption. (Huber et al., 2019). These studies, and others like them, demonstrate indirect links to energy derivation in neuronal and adipocyte mitochondria, but none show a direct energy-derivation or facilitation role for NAA. Our model is the first attempt to demonstrate a biochemically meaningful thermodynamic effect exerted by ASPNAT on a core mitochondrial metabolic node.

### 4.9 Similarities between ASPNAT, Aralar1, ASPA and GOT2 deficiencies

Two enzymes and one transporter modeled here, ASPNAT, GOT2 and Aralar1, lie at a crossroads of mitochondrial metabolism, where the malate-aspartate shuttle interfaces with the TCA cycle. Bottlenecks at this juncture are highly detrimental to mitochondrial function and defects in these and related proteins can have similar effects on neurological development. Further, NAA metabolism requires the enzyme aspartoacylase (ASPA) which hydrolyzes NAA into acetate and aspartate, and therefore, ASPA deficiency can provide information on the importance of NAA catabolism.

ASPNAT deficiency in mice results in behavioral abnormalities (Ariyannur et al., 2013; Furukawa-Hibi et al., 2012; Sumi et al., 2017; Toriumi et al., 2018), possibly due to disrupted neurogenesis and altered neuronal morphology (Wulaer et al., 2021). ASPNAT deficient mice raised on a fat-free diet die before reaching maturity, and NAA supplementation of the fat-free diet rescues the developing mice (Hofer et al., 2019). In humans, ASPNAT deficiency produces a developmental disorder with microcephaly, hypomyelination and seizures (Boltshauser et al., 2004).

ASPA deficiency results in a well-characterized leukodystrophy known as Canavan disease (CD). CD leads to hypomyelination, hypotonia, arrested psychomotor development, brain vacuolization and seizures (Matalon et al., 1995; Matalon et al., 1988). ASPA deficiency in mice raises blood NAA levels, which in turn directs glucose handling toward pyrimidine synthesis in adipose tissue. This effect was shown to be due to the elevated circulating NAA levels (Felix et al., 2025). Cytoplasmic aspartate is an obligate substrate for pyrimidine synthesis, and NAA hydrolysis by ASPA provides additional aspartate for this metabolic pathway. These findings implicate NAA in glucose handling, and show that NAA shifts metabolism toward anabolic activities like increased pyrimidine synthesis (Felix et al., 2025). While focus on NAA metabolism has highlighted acetate delivery to the cytoplasm for lipid synthesis, and the cell nucleus for histone acetylation, it now seems likely that NAA export from mitochondria also provides additional cytoplasmic aspartate to support anabolic processes, including pyrimidine, purine, asparagine, arginine, NAD^+^ and protein synthesis, among others.

In humans, Aralar1 (*AGC1*) deficiency results in global hypomyelination, severe hypotonia, arrested psychomotor development, and seizures beginning at a few months of age (Wibom et al., 2009). GOT2 deficiency leads to progressive neurodevelopmental delay, severe to profound intellectual disability, infantile epilepsy, progressive microcephaly, and hypotonia evolving into spasticity with axial hypotonia (German et al., 2025). Low aspartate was also observed in some GOT2 deficient patients.

Common features to ASPA, GOT2 and Aralar1 deficiencies include developmental delay or arrest, hypomyelination, hypotonia, ataxia and seizures. ASPNAT deficiency in humans also results in hypomyelination and seizures. These neurodevelopmental outcomes result, at least in part, from defects in mitochondrial energy derivation or cytoplasmic acetyl-CoA metabolism in the brain during development. We have shown that NAA synthesis by ASPNAT provides a modest but significant facilitation of AAT throughput, with the degree of the effect being largely dependent upon the concentrations of the substrates and products. The results support the conclusion that ASPNAT can serve as an auxiliary matrix-aspartate sink when the AAT node is product-limited.

## 5. Conclusions

In the current study, we have shown that ASPNAT facilitates mitochondrial metabolism through multiple mechanisms including removal of product inhibition for AAT, enhancement of AAT throughput, especially at low oxaloacetate concentration regimes, and facilitation of glutamate oxidation in the TCA cycle. This parallel pathway to citrate synthase for carbon transport from the mitochondrial matrix to the cytoplasm clearly provides metabolic advantages as ASPNAT is phylogenetically conserved among vertebrates. Adipocytes utilize the same metabolic mechanism within one cell type, whereas in the nervous system, the NAA synthesized by ASPNAT must first be transported to oligodendrocytes before being reconverted to acetyl-CoA for lipid synthesis in oligodendrocytes. The two-cell system for NAA synthesis and utilization in the nervous system is similar to other such systems, such as glutamate-glutamine cycling between neurons and astrocytes, where the metabolic output of one cell type is a required input for the other.

The present modeling of the mitochondrial aspartate node helps clarify ASPNAT’s roles in specialized acetyl-CoA metabolism in the nervous system and is potentially extendable to adipocytes. Biological systems are replete with redundancy, parallel pathways and pathway reversibility, and it is instructive to view NAA synthesis in this light as allowing for an additional method for removing aspartate from the mitochondrial matrix, while also providing a secondary pathway to cytoplasmic acetyl-CoA synthesis. This alternate pathway to cytoplasmic acetyl-CoA synthesis is expressed in cell types with high rates of lipid synthesis and turnover, including the nervous system and adipose tissue, and it becomes more important when mitochondrial citrate synthase is saturated.

These results could be tested experimentally. Several predictions can be made about ASPNAT inhibition in cell culture with small molecule inhibitors (Nesuta et al., 2021) or ASPNAT knockout. Loss of ASPNAT function should decrease mitochondrial aspartate levels, increase the aspartate/αKG ratio and increase aspartate export from mitochondria. Declining NAA synthesis would be accompanied by impaired GOT2/MAS function. ASPNAT functional loss should reduce mitochondrial production of αKG, confirming the bottleneck at GOT2/AAT. These could potentially be tested using isotope-tracing, mass-labeling, mitochondrial metabolomics and ASPNAT inhibitors or knockout in cell culture or with isolated brain mitochondrial preparations.

## 6 Limitations

The current study is focused on the thermodynamic effects of NAA synthesis on mitochondrial AAT throughput. This restricted model employed clamped boundary pools, a semi-empirical effective ASPNAT capacity and estimated mitochondrial OAA and acetyl-CoA setpoints. Many other mitochondrial and cytosolic effects that incur from NAA synthesis, or lack thereof, have not been considered here. It should be noted that citrate synthesis is a factor that was not modeled here, but it would be expected that when citrate synthase was near saturation, NAA synthesis would be increased, facilitating AAT throughput. Another limitation is the lack of a dicarboxylate transporter that moves NAA to the cytoplasm. To date, the mitochondrial NAA exporter has not been identified and, as such, could not be accurately included in the model. Therefore, the model employs ASPA action as a substitute for NAA clearance. If the NAA transporter is not a major restriction for NAA export, then ASPA action would be the primary site of NAA clearance. The model also does not take into account other boundary conditions, such as the supply of malate via the oxoglutarate carrier, which is needed for aspartate synthesis by AAT. The model is therefore best viewed as a constrained mechanistic system that predicts signs, dependencies, and regime boundaries more confidently than absolute whole-neuron or adipocyte metabolite fluxes. Future studies will incorporate additional relevant enzymes and transporters involved in the aspartate node of mitochondrial metabolism to improve the model’s accuracy. Expanded mathematical details, robustness analyses, and parameter-status tables are provided in Supplementary Methods and **Supplementary Tables S10 to S20**.

## 7 Author Contributions

NP, JRM, BSS, and AMN conceived the study. NP developed the computational model and performed the simulations. NP, JRM, BSS, and AMN interpreted the results and wrote or edited the manuscript. AMN and BSS provided funding. All authors approved the submitted version.

## Data Availability Statement

All data and models generated are provided in the supplementary information. No new software was generated in this study.

## Disclaimer

The opinions and assertions expressed herein are those of the authors and do not necessarily reflect the official policy or position of the Uniformed Services University or the DoW.

