## Supplementary material for "N-Acetylaspartate Synthesis as a Thermodynamic Relief Mechanism for Mitochondrial Aspartate Aminotransferase"

#### 1 1. Supplementary Methods

This document contains the full rate-law specification, the full Antimony implementation, the extended parameter and calibration tables, and the full-size robustness figures referenced by the main manuscript. The supplementary material is organized to complement rather than restate the main text. The emphasis is on model definition, parameter provenance, and the additional numerical checks that constrain interpretation of the relief-valve mechanism.

The reduced model is presented in the standard systems-biology sense of a scoped biochemical network translated into dynamic state variables, reaction fluxes, boundary conditions, and mass-balance equations (Sauro and Kholodenko 2004; Sauro and Bergmann 2010; Alon 2006; Palsson 2006; Jamshidi and Palsson 2010). These references support the modeling grammar and dynamical framework only. They are not used here to import genome-scale flux-balance assumptions, transcriptional circuit motifs, or whole-cell reconstruction claims into this localized mitochondrial node.

##### 1.1 Full rate law specification

The reduced mitochondrial node contains three dynamic species, matrix aspartate, matrix $\alpha$-ketoglutarate, and matrix N-acetylaspartate (NAA). Glutamate, oxaloacetate (OAA), acetyl-CoA, and coenzyme A (CoA) are treated as clamped boundary pools so that the model isolates the thermodynamically sensitive mitochondrial aspartate node while retaining the dominant chemical drivers of the reaction system. Aspartate aminotransferase (AAT) constants were taken from synaptic mitochondrial preparations, and the reverse AAT capacity was constrained by detailed balance using the equilibrium relation measured in isotope-exchange studies (McKenna et al. 2006; Kimmich, Roussie, and Randles 2002). ASPNAT substrate and inhibition terms were anchored to detergent-solubilized enzymology measurements and interpreted alongside later stabilization studies that support higher effective activity in native membrane-associated settings (Madhavarao et al. 2003; Wang et al. 2016).

The AAT rate law was

$$v_{\mathrm{AAT}}=\frac{\frac{V_f[\mathrm{Glu}][\mathrm{OAA}]}{K_{\mathrm{Glu}}K_{\mathrm{OAA}}}-\frac{V_r[\mathrm{Asp}][\alpha\mathrm{KG}]}{K_{\mathrm{Asp}}K_{\alpha\mathrm{KG}}}}{\left(1+\frac{[\mathrm{Glu}]}{K_{\mathrm{Glu}}}+\frac{[\mathrm{Asp}]}{K_{\mathrm{Asp}}}\right)\left(1+\frac{[\mathrm{OAA}]}{K_{\mathrm{OAA}}}+\frac{[\alpha\mathrm{KG}]}{K_{\alpha\mathrm{KG}}}\right)},$$

with the reverse capacity constrained by

$$V_r=\frac{V_f K_{\alpha\mathrm{KG}}K_{\mathrm{Asp}}}{K_{\mathrm{eq}}K_{\mathrm{Glu}}K_{\mathrm{OAA}}}.$$

ASPNAT was represented as

$$v_{\mathrm{ASPNAT}}=V_{\mathrm{ASPNAT}}\cdot\frac{[\mathrm{Asp}]}{K_{\mathrm{Asp}}^{\mathrm{ASPNAT}}+[\mathrm{Asp}]}\cdot\frac{[\mathrm{AcCoA}]}{K_{\mathrm{AcCoA}}+[\mathrm{AcCoA}]}\cdot\frac{1}{1+[\mathrm{NAA}]/\mathrm{IC}_{50}^{\mathrm{NAA}}}\cdot\frac{1}{1+[\mathrm{CoA}]/\mathrm{IC}_{50}^{\mathrm{CoA}}}.$$

Baseline AGC1-linked aspartate removal was approximated by a first-order term, $v_{\mathrm{export}}=k_{\mathrm{export}}[\mathrm{Asp}]$, and a Michaelis-Menten replacement was tested separately as a robustness check. The state equations follow directly from these rates

$$\begin{aligned}\frac{d[\mathrm{Asp}]}{dt}&=v_{\mathrm{AAT}}-v_{\mathrm{ASPNAT}}-v_{\mathrm{export}},\\ \frac{d[\alpha\mathrm{KG}]}{dt}&=v_{\mathrm{AAT}}-k_{\alpha\mathrm{KG}}[\alpha\mathrm{KG}],\\ \frac{d[\mathrm{NAA}]}{dt}&=v_{\mathrm{ASPNAT}}-k_{\mathrm{NAA}}[\mathrm{NAA}],\end{aligned}$$

and the near-equilibrium reversal boundary is defined by $[\mathrm{Asp}][\alpha\mathrm{KG}]/([\mathrm{Glu}][\mathrm{OAA}])=K_{\mathrm{eq}}$.

#### 1.2 Calibration and numerical workflow

The effective ASPNAT capacity was treated as semi-empirical because detergent-solubilized preparations substantially under-report the activity expected for the native enzyme in membrane-associated neuronal compartments (Madhavarao et al. 2003; Wang et al. 2016). The calibrated value used in the baseline model was chosen to remain consistent with reported NAA synthesis and steady-state NAA levels while preserving physically plausible substrate and inhibition terms. The main claims do not depend on one exact ASPNAT value. They were tested explicitly across broad capacity sweeps and across the global uncertainty designs reported below.

All steady states were obtained with relative tolerance $10^{-10}$ and absolute tolerance $10^{-13}$ (Petzold 1983). The baseline workflow used the Livermore Solver for Ordinary Differential Equations with Automatic method switching (LSODA) and extended each integration in 1000 min blocks until the maximum absolute derivative across the dynamic species fell below $10^{-9}$. If the dynamic-OAA extension failed that residual test under LSODA, the same branch was re-integrated with a stiff solver until the same derivative criterion was met. Global uncertainty analysis used Latin Hypercube Sampling across ten uncertain parameters with $N=5000$ samples. A separate correlated-sampling stress test asked whether strong OAA, acetyl-CoA, and export-related covariance structures altered the direction of the relief effect. The two-compartment variant is therefore interpreted as a topological sign test, whereas the dynamic-OAA variant is interpreted as a solver-checked sensitivity analysis rather than as an independent physiological benchmark.

##### 1.3 Full Antimony model

The following Antimony listing gives the full baseline model in a human-readable form that can be converted to SBML or simulated with standard systems-biology tools (Smith et al. 2009; Hucka et al. 2003; K. Choi et al. 2018; Medley et al. 2018; Porubsky and Sauro 2023). Antimony is cited here as the text-based model specification language, SBML as the model-exchange standard, and Tellurium-related work as compatible reproducible dynamical-modeling infrastructure. The present supplement provides the mathematical specification and Antimony listing, not a full SED-ML or COMBINE archive.

```
model NAA_ReliefValve()

    // --- Compartment ---
    compartment mito = 1.0;  // normalized

    // --- Dynamic Species ---
    species Asp in mito;   // mitochondrial aspartate (mM)
    species aKG in mito;   // alpha-ketoglutarate (mM)
    species NAA in mito;   // N-acetylaspartate (mM)

    // --- Clamped Boundary Species ---
    const species Glu in mito;   // glutamate (mM)
    const species OAA in mito;   // oxaloacetate (mM)
    const species AcCoA in mito; // acetyl-CoA (mM)
    const species CoASH in mito; // coenzyme A (mM)

    // --- Initial Conditions ---
    Asp   = 0.050;    // 50 uM
    aKG   = 0.050;    // 50 uM
    NAA   = 0.010;    // 10 uM
    Glu   = 10.0;     // 10 mM (clamped)
    OAA   = 0.000160; // 160 nM (clamped)
    AcCoA = 0.014;    // 14 uM (clamped)
    CoASH = 0.026;    // 26 uM (clamped)

    // ============================================================
    // REACTION 1: AAT (GOT2) -- Reversible Ping-Pong Bi-Bi
    // Glu + OAA <-> Asp + aKG
    // ============================================================
    Vr_AAT := Vf_AAT * Km_aKG * Km_Asp_AAT /
              (Keq_AAT * Km_Glu * Km_OAA);

    J_AAT: Glu + OAA -> Asp + aKG;
        (Vf_AAT * Glu * OAA / (Km_Glu * Km_OAA)
         - Vr_AAT * Asp * aKG / (Km_aKG * Km_Asp_AAT)) /
```

```
    ((1 + Glu/Km_Glu + Asp/Km_Asp_AAT) *
     (1 + OAA/Km_OAA + aKG/Km_aKG));

// ===========================================================
// REACTION 2: ASPNAT -- Irreversible Bi-Substrate
// Asp + AcCoA -> NAA + CoA
// ===========================================================
J_ASPNAT: Asp + AcCoA -> NAA + CoASH;
    Vmax_ASPNAT
    * (Asp / (Km_Asp_ASPNAT + Asp))
    * (AcCoA / (Km_AcCoA + AcCoA))
    * (1 / (1 + NAA / IC50_NAA))
    * (1 / (1 + CoASH / IC50_CoA));

// ===========================================================
// REACTION 3: Aspartate Export
// ===========================================================
J_export: Asp -> ; k_export * Asp;

// ===========================================================
// REACTION 4: alpha-KG Removal
// ===========================================================
J_aKG_removal: aKG -> ; k_aKG * aKG;

// ===========================================================
// REACTION 5: NAA Removal
// ===========================================================
J_NAA_export: NAA -> ; k_NAA * NAA;

// ===========================================================
// KINETIC PARAMETERS
// ===========================================================
Vf_AAT      = 1615.0;
Km_Glu      = 8.9;
Km_OAA      = 0.050;
Km_Asp_AAT  = 2.0;
Km_aKG      = 0.64;
Keq_AAT     = 6.6;

Vmax_ASPNAT   = 29.0;
Km_Asp_ASPNAT = 0.580;
Km_AcCoA     = 0.058;
IC50_NAA     = 0.850;
IC50_CoA     = 0.420;

k_export = 1.0;
k_aKG    = 2.0;
```

```
    k_NAA    = 0.01;

end
```

### 2 Supplementary Tables

#### 2.1 Parameter architecture and provenance

The parameter program combines direct biochemical measurements, thermodynamic constraints, tissue-scale metabolite estimates, and effective rate constants chosen to represent unresolved transport and turnover processes. The core AAT values are anchored in synaptic mitochondrial measurements (McKenna et al. 2006). ASPNAT substrate constants and inhibition terms come from the original enzyme characterization (Madhavarao et al. 2003), with the effective activity interpreted in light of later recombinant stabilization data (Wang et al. 2016). Boundary metabolite pools were chosen to represent the low-OAA neuronal regime discussed in the main text and the acetyl-group buffering ranges reported for brain mitochondria (Erecińska et al. 1988; Ronowska et al. 2018).

**Table S1:** *Core parameter set used for the reduced mitochondrial-node model. This table consolidates the complete parameter architecture, provenance notes, and boundary-pool definitions used in the supplementary analyses.*

| **Parameter** | **Value** | **Units** | **Provenance and note** |
|---|---|---|---|
| **Parameter** | **Value** | **Units** | **Provenance and note** |
| $V_f$ (AAT forward capacity) | 1615 | nmol/min/mg | Synaptic mitochondrial AAT activity (McKenna et al. 2006). |
| $V_r$ (Haldane) | 703.8 | nmol/min/mg | Calculated from the detailed-balance relation with $K_{\text{eq}} = 6.6$. |
| $K_{\text{Glu}}$ | 8.9 | mM | AAT substrate constant (McKenna et al. 2006). |
| $K_{\text{OAA}}$ | 0.050 | mM | Literature-supported mid-range value for mitochondrial AAT. |
| $K_{\text{Asp}}$ | 2.0 | mM | AAT product constant (McKenna et al. 2006). |
| $K_{\alpha\text{KG}}$ | 0.64 | mM | High-affinity synaptic mitochondrial branch (McKenna et al. 2006). |
| $K_{\text{eq}}$ | 6.6 | n/a | AAT equilibrium constant from isotope-exchange measurements (Kimmich, Roussie, and Randles 2002). |
| $V_{\text{ASPNAT}}$ | 29 | nmol/min/mg | Semi-empirical effective capacity chosen to match observed NAA synthesis within a biologically credible range. |
| $K_{\text{Asp}}^{\text{ASPNAT}}$ | 0.580 | mM | ASPNAT substrate constant (Madhavarao et al. 2003). |
| $K_{\text{AcCoA}}$ | 0.058 | mM | ASPNAT acetyl-CoA constant (Madhavarao et al. 2003). |

| Parameter | Value | Units | Provenance and note |
|---|---|---|---|
| $\mathrm{IC}_{50}^{\mathrm{NAA}}$ | 0.850 | mM | ASPNAT product inhibition by NAA (Madhavarao et al. 2003). |
| $\mathrm{IC}_{50}^{\mathrm{CoA}}$ | 0.420 | mM | ASPNAT product inhibition by CoA (Madhavarao et al. 2003). |
| $[\mathrm{Glu}]$ | 10.0 | mM | Estimated neuronal mitochondrial glutamate pool (Erecińska et al. 1988). |
| $[\mathrm{OAA}]$ | $1.6 \times 10^{-4}$ | mM | Baseline clamped OAA that represents the low-OAA neuronal regime. |
| $[\mathrm{AcCoA}]$ | 0.014 | mM | Brain mitochondrial acetyl-CoA estimate (Ronowska et al. 2018). |
| $[\mathrm{CoA}]$ | 0.026 | mM | Brain mitochondrial coenzyme A estimate (Ronowska et al. 2018). |
| $k_{\mathrm{export}}$ | 1.0 | $\mathrm{min}^{-1}$ | Effective AGC1-linked aspartate export in the baseline model. |
| $k_{\alpha\mathrm{KG}}$ | 2.0 | $\mathrm{min}^{-1}$ | Effective downstream $\alpha$-ketoglutarate removal. |
| $k_{\mathrm{NAA}}$ | 0.01 | $\mathrm{min}^{-1}$ | Effective matrix NAA removal with $\tau = 100$ min. |
| Matrix volume | 1.0 | $\mu$L/mg protein | Conventional conversion used to map nmol/min/mg to mM/min. |
| Protein per tissue | 100 | mg protein/g | Approximate brain homogenate conversion factor. |

**Table S2:** *S2AAT kinetic parameters in synaptic and non-synaptic mitochondria.*

| Parameter | Synaptic | Non-synaptic | Unit |
|---|---|---|---|
| $K_m^{\alpha\mathrm{KG}}$ (total) | 1.95 | 1.47 | mM |
| $K_m^{\alpha\mathrm{KG}}$ (high affinity) | 0.64 | 0.46 | mM |
| $K_m^{\alpha\mathrm{KG}}$ (low affinity) | 3.92 | 3.08 | mM |
| $V_{\mathrm{max}}$ (total) | 1615 | 1842 | nmol/min/mg |
| $V_{\mathrm{max}}$ (high affinity) | 689 | 734 | nmol/min/mg |
| $V_{\mathrm{max}}$ (low affinity) | 1947 | 2620 | nmol/min/mg |

Source table adapted from McKenna et al. (2006). The model uses the synaptic high-affinity branch because synaptic mitochondria dominate rapid neuronal glutamate handling.

#### 2.2 Steady-state, energetic, and validation outputs

The baseline steady state supports the current claim that ASPNAT lowers matrix aspartate while increasing forward AAT throughput. The energetic tables are included separately because they are useful for auditing scale-dependent interpretations. The validation table is retained to show which outputs are directly anchored to independent measurements and which remain model-level predictions.

**Table S3:** *Steady-state concentrations and fluxes with ASPNAT active or inactive in the baseline model.*

| Variable | ASPNAT on | ASPNAT off | Change | Unit |
|---|---|---|---|---|
| *Concentrations* | | | | |
| [Asp] | 105.37 | 140.95 | $+33.8\%$ | $\mu$M |
| [$\alpha$KG] | 92.27 | 70.48 | $-23.6\%$ | $\mu$M |
| [NAA] | 7.916 | 0.000 | $-100.0\%$ | mM |
| *Fluxes* | | | | |
| $v_{\text{AAT}}$ | 0.1845 | 0.1410 | $-23.6\%$ | mM/min† |
| $v_{\text{ASPNAT}}$ | 0.0792 | 0.0000 | $-100.0\%$ | mM/min† |
| $v_{\text{Asp export}}$ | 0.1054 | 0.1410 | $+33.8\%$ | mM/min† |
| Total Asp removal | 0.1845 | 0.1410 | $-23.6\%$ | mM/min† |
| *Thermodynamic indicators* | | | | |
| $\Gamma$ | 6.08 | 6.17 | $+1.5\%$ | n/a |
| $\Gamma/K_{\text{eq}}$ | 0.921 | 0.935 | $+1.5\%$ | n/a |
| $\Delta G$ | $-0.21$ | $-0.17$ | n/a | kJ/mol |
| Asp threshold for forward AAT | 114.5 | 149.8 | n/a | $\mu$M |

†Model flux units are numerically equal to nmol/min/mg under the $V_{\text{eff}} = 1$ $\mu$L/mg convention used throughout the manuscript.

**Table S4:** *Comparative steady-state proxy indices for the local mitochondrial node at baseline conditions. The relative differences are robust. The absolute values are local proxy quantities generated by a reduced, clamped micro-domain model and should not be scaled into whole-cell energetic budgets.*

| Index | ASPNAT on | ASPNAT off | $\Delta$ | Relative $\Delta$ | Unit |
|---|---|---|---|---|---|
| MAS support fraction | 1.27 | 0.97 | $+0.30$ | $+30.9\%$ | % |
| ATP from AAT† | 1.845 | 1.410 | $+0.436$ | $+31\%$ | mfu |
| ATP fraction of total | 0.739 | 0.565 | $+0.175$ | $+30.9\%$ | % |
| AcCoA drain by ASPNAT | 0.507 | 0.00 | $+0.507$ | n/a | % of PDH |
| AcCoA/CoA effective ratio | 0.534 | 0.539 | $-0.004$ | $-0.8\%$ | ratio |

**Table S5:** *Scaled sensitivity coefficients for all fitted or estimated parameters.*

| Parameter | $S_{AAT}$ | $S_{ASPNAT}$ | Interpretation |
|---|---|---|---|
| *AAT kinetic parameters* | | | |
| $V_f$ (AAT) | $+0.035$ | $+0.021$ | Near-zero local control in the baseline state |
| $K_{Glu}$ | $-0.017$ | $-0.010$ | Glutamate is effectively saturating |
| $K_{OAA}$ | $-0.036$ | $-0.021$ | OAA control remains modest locally |
| $K_{Asp}$ | $+0.001$ | $+0.001$ | Asp product term is negligible locally |
| $K_{\alpha KG}$ | $+0.004$ | $+0.003$ | $\alpha$KG product term is minor |
| $K_{eq}$ | $+0.413$ | $+0.241$ | Dominant thermodynamic driver |
| *ASPNAT kinetic parameters* | | | |
| $V_{ASPNAT}$ | $+0.123$ | $+0.464$ | Strong self-control of ASPNAT flux |
| $K_m^{ASPNAT}$ Asp | $-0.103$ | $-0.392$ | Asp affinity strongly controls ASPNAT flux |
| $K_m^{ASPNAT}$ AcCoA | $-0.098$ | $-0.373$ | AcCoA affinity strongly controls ASPNAT flux |
| $IC_{50}^{NAA}$ | $+0.111$ | $+0.419$ | NAA feedback is a major regulator |
| $IC_{50}^{CoA}$ | $+0.007$ | $+0.027$ | CoA feedback is minor |
| *Boundary species* | | | |
| [Glu] | $+0.431$ | $+0.251$ | Dominant substrate supply term |
| [OAA] | $+0.449$ | $+0.262$ | Dominant thermodynamic driver |
| [AcCoA] | $+0.099$ | $+0.373$ | Metabolic gate on ASPNAT |
| *Removal rates* | | | |
| $k_{export}$ | $+0.311$ | $-0.151$ | Competing product-removal branch |
| $k_{\alpha KG}$ | $+0.418$ | $+0.244$ | Major AAT support term |
| $k_{NAA}$ | $+0.111$ | $+0.419$ | Moderate control through NAA removal |

The opposite signs for $k_{export}$ in the AAT and ASPNAT columns quantify the competition between AGC1-mediated export and ASPNAT-mediated product removal.

**Table S6:** *Comparison of model outputs with independent experimental measurements.*

| Observable | Model | Experimental | Source |
|---|---|---|---|
| NAA synthesis rate | nmol/min/mg (≈7.9 ) | in human and 10 nmol $g^{-1}$ 12 nmol $g^{--1}$ in rat | (Moreno, Ross, and Blüml 2001; Truckenmiller et al. 1985) |
| Steady-state NAA | mM | 8 –14 in whole tissue and about 20 mM intracellularly | (Moffett et al. 2006; Moffett et al. 2007) |

| Observable | Model | Experimental | Source |
|---|---|---|---|
| Matrix aspartate | µM | 2 –4 total neuronal aspartate | (Erecińska et al. 1988) |
| Matrix NAA removal | $\tau = 100min$ and $t_{1/2} \approx 69min$ | Whole-brain turnover on the order of 16 h–70 h | (I.-Y. Choi and Gruetter 2004) |
| AAT near-equilibrium state | $\Gamma/K_{\mathrm{eq}} = 0.92$ | Extensive isotope exchange consistent with near-equilibrium behavior | (Kimmich, Roussie, and Randles 2002) |

The matrix-compartment predictions should not be compared numerically with whole-cell pools without regard to compartment size. The comparison is intended to show directional and order-of-magnitude consistency rather than a literal one-to-one mapping.

**Table S7:** *ASPNAT activity measurements across preparations.*

| Source | Preparation | Activity (nmol/min/mg) | Note |
|---|---|---|---|
| Madhavarao et al. 2003 | CHAPS-solubilized rat brain | 0.45 | Detergent-solubilized preparation with marked activity loss (Madhavarao et al. 2003). |
| Madhavarao 2003 corrected | Estimated native activity | ≥9.0 | Back-calculated from the reported >95% activity loss after CHAPS treatment. |
| This model | Effective tissue-level capacity | 29.0 | Semi-empirical value required to match steady-state NAA synthesis in the low-substrate regime. |
| Wang et al. 2016 | MBP-ANAT fusion protein | 70.0 | Stabilized recombinant human ASPNAT with preserved catalytic capacity (Wang et al. 2016). |

**Table S8:** *Energetic conversion factors used in stoichiometric projections.*

| Parameter | Value | Unit | Source and justification |
|---|---|---|---|
| **Parameter** | **Value** | **Unit** | **Source and justification** |
| *Oxidative phosphorylation P/O ratios* | | | |
| P/O for NADH | 2.5 | ATP/NADH | Modern consensus summarized by Dienel (2019). |
| P/O for $FADH_2$ | 1.5 | ATP/$FADH_2$ | Modern consensus summarized by Dienel (2019). |
| ATP per glucose | 32 | ATP/glucose | Standard oxidative accounting. |
| *TCA segment from $\alpha$KG to OAA* | | | |
| $\alpha$KG dehydrogenase | 2.5 | ATP equivalent | 1 NADH times 2.5. |

| Parameter | Value | Unit | Source and justification |
|---|---|---|---|
| Succinyl-CoA synthetase | 1.0 | ATP equivalent | 1 GTP. |
| Succinate dehydrogenase | 1.5 | ATP equivalent | 1 $FADH_2$ times 1.5. |
| Malate dehydrogenase | 2.5 | ATP equivalent | 1 NADH times 2.5. |
| Total from $\alpha$KG to OAA | 7.5 | ATP equivalent | Sum of the segmental terms. |
| *MAS shuttle* | | | |
| NADH per MAS cycle | 1.0 | NADH | One reducing equivalent per cycle. |
| ATP per MAS-derived NADH | 2.5 | ATP | Entry at Complex I. |
| *Combined ATP per AAT turnover* | | | |
| MAS component | 2.5 | ATP | Shuttle-linked reducing equivalent. |
| TCA component | 7.5 | ATP | Oxidation of $\alpha$KG back to OAA. |
| Total per AAT turnover | 10.0 | ATP | Combined yield. |
| *Unit conversions* | | | |
| $CMR_{glc}$ in neurons | 0.73 | $\mu$mol/g/min | Dienel (2019). |
| Protein per tissue | 100 | mg protein/g | Approximate brain conversion. |
| Baseline glycolytic NADH | 14.6 | nmol/min/mg | Derived from glucose consumption. |
| Baseline MAS fraction | 93 | % | Dienel (2019). |
| Baseline MAS flux | 13.6 | nmol/min/mg | Derived from the glycolytic NADH term. |
| Total neuronal ATP | $\sim$250 | nmol/min/mg | Approximate oxidative budget. |
| PDH flux | 14.6 | nmol/min/mg | Equal to $2 \times \mathrm{CMR_{glc}} \times 10$. |

**Table S9:** *Summary statistics of the two-dimensional relief map evaluated on a $25 \times 25$ grid. The full grid includes extreme low-OAA, low-export states, so the upper tail is intentionally skewed and the maximum should not be read as a representative physiological operating point.*

| Statistic | Value |
|---|---|
| Grid dimensions | $25 \times 25$ |
| OAA range | 10 –3162 |
| $k_{\mathrm{export}}$ range | 0.05 min–3.0 min |
| Maximum relief | 407.5% |
| Minimum relief | 2.71% |
| Mean relief | 26.94% |

| Statistic | Value |
|---|---|
| Median relief | 14.02% |
| *Physiological reference point* | |
| OAA of 160 nM and $k_{\text{export}} = 1.0$ | about 30.9% |
| *Monotonicity* | |
| Relief versus OAA at fixed export | monotonically decreasing |
| Relief versus export at fixed OAA | monotonically decreasing |

**Table S10:** *MAS support fraction, ATP from AAT, acetyl-CoA drain, and effective acetyl-CoA to coenzyme A ratio across the ASPNAT titration sweep.*

| ASPNAT (%) | MAS (%) | ATP from AAT (nmol/min/mg) | AcCoA drain (%) (of PDH) | AcCoA/CoA (effective) |
|---|---|---|---|---|
| **ASPNAT (%)** | **MAS (%)** | **ATP from AAT** (nmol/min/mg) | **AcCoA drain (%)** (of PDH) | **AcCoA/CoA** (effective) |
| | 0.97 | 1.410 | 0.000 | 0.5385 |
| 25.4 | 1.12 | 1.620 | 0.260 | 0.5362 |
| 50.8 | 1.18 | 1.715 | 0.368 | 0.5352 |
| 76.3 | 1.23 | 1.788 | 0.447 | 0.5345 |
| 101.7 | 1.27 | 1.849 | 0.511 | 0.5339 |
| 127.1 | 1.31 | 1.903 | 0.567 | 0.5335 |
| 147.5 | 1.34 | 1.942 | 0.606 | 0.5331 |
| 172.9 | 1.37 | 1.987 | 0.651 | 0.5327 |
| 198.3 | 1.40 | 2.029 | 0.692 | 0.5324 |
| 223.7 | 1.43 | 2.068 | 0.730 | 0.5320 |
| 249.2 | 1.45 | 2.105 | 0.765 | 0.5317 |
| 274.6 | 1.47 | 2.140 | 0.798 | 0.5314 |
| 300.0 | 1.50 | 2.173 | 0.829 | 0.5312 |

**Table S11:** *Complete energetic comparison between ASPNAT-active and ASPNAT-inactive steady states.*

| Index | On | Off | $\Delta$ |
|---|---|---|---|
| Cytoplasmic $NAD^+$/NADH | 1000.13 | 1000.10 | $+0.03$ |
| MAS support fraction (%) | 1.27 | 0.97 | $+0.30$ |
| Mitochondrial NADH flux$^{\dagger}$ | 0.369 | 0.282 | $+0.087$ |
| ATP from AAT$^{\dagger}$ | 1.845 | 1.410 | $+0.436$ |

| Index | On | Off | $\Delta$ |
|---|---|---|---|
| ATP fraction of total (%) | 0.739 | 0.565 | $+0.175$ |
| MAS-dependent ATP (%) | 0.185 | 0.141 | $+0.044$ |
| AcCoA drain by ASPNAT (% of PDH) | 0.507 | 0.000 | $+0.507$ |
| AcCoA/CoA effective ratio | 0.534 | 0.539 | $-0.004$ |
| AcCoA fraction versus citrate synthase (%) | 0.543 | 0.000 | $+0.543$ |

†These values are scale-dependent absolute magnitudes. The robust conclusion is the direction of the tradeoff. The energetic gain associated with enhanced AAT flux is modest and is partly offset by the acetyl-CoA cost of NAA synthesis.

The ensemble was intentionally broad rather than fitted sample-by-sample to reproduce a single baseline relief value. Parameter draws were required only to remain within the biochemical and boundary-condition ranges in Table ; no post hoc constraint forced a target AAT-relief percentage or a target NAA concentration.

**Table S12:** *Parameter distributions used for the global uncertainty analysis with* $N = 5000$ *samples.*

| Parameter | Distribution | Range | Rationale |
|---|---|---|---|
| $[\mathrm{OAA}]$ | LogUniform | 50 to 500 nM | Ischemia to mild activation |
| $[\mathrm{AcCoA}]$ | LogUniform | 5 to 50 $\mu$M | Dietary and metabolic range |
| $k_{\text{export}}$ | Uniform | 0.3 to 3.0 $\text{min}^{-1}$ | AGC1 capacity range |
| $k_{\alpha\text{KG}}$ | Uniform | 1.0 to 4.0 $\text{min}^{-1}$ | $\alpha$KGDH activity range |
| $V_{\text{ASPNAT}}$ | LogUniform | 5 to 100 nmol/min/mg | Wide plausible capacity range |
| $\mathrm{IC}_{50}^{\text{NAA}}$ | Uniform | 0.5 to 1.5 mM | Literature uncertainty |
| $\mathrm{IC}_{50}^{\text{CoA}}$ | Uniform | 0.2 to 0.8 mM | Literature uncertainty |
| $K_{\text{eq}}$ | Uniform | 5.0 to 8.0 | Thermodynamic uncertainty |
| $K_m^{\text{OAA}}$ | Uniform | 0.030 to 0.100 mM | Measurement range |
| $k_{\text{NAA}}$ | LogUniform | 0.005 to 0.02 $\text{min}^{-1}$ | Matrix-turnover range |

**Table S13:** *Relief comparison for first-order and Michaelis-Menten AGC1 kinetics.*

| Transport model | $\mathbf{K_m}$ (mM) | $\mathbf{V_{max}}$ (mM/min) | Relief (%) |
|---|---|---|---|
| First-order baseline | n/a | n/a | 30.9 |
| Michaelis-Menten | 0.5 | 0.605 | 35.1 |
| Michaelis-Menten | 2.0 | 2.105 | 32.0 |

#### 2.3 Dynamic and topology support

The next table cluster records the numerical checks that constrain how far the current mechanism can be interpreted. The dynamic-OAA variant asks whether endogenous OAA turnover changes the sign of the effect, but it is reported only as a solver-checked sensitivity analysis because the ASPNAT-off branch is stiff. The two-compartment variant asks whether stress-state rerouting of effective aspartate consumption would change the sign of the effect; it is not treated as evidence against mitochondrial ASPNAT as the default biological premise. The remaining tables document the flux scale implied by the effective rate constants and the uncertainty classes assigned to the global sampling program.

**Table S14:** *Parameters used in the dynamic-OAA sensitivity extension. The MDH2 and citrate-synthase terms were set to place this reduced robustness check in the same low-OAA regime as the clamped reference model; they are not a neuronal whole-mitochondrion calibration.*

| Parameter | Value | Unit | Source |
|---|---|---|---|
| $V_f^{\mathrm{MDH}}$ | 50 | mM/min | Calibrated to place the extension in the same low-OAA regime as the baseline model |
| $K_{\mathrm{eq}}^{\mathrm{MDH}}$ | $10^{-5}$ | n/a | Strongly favors malate |
| $K_m^{\mathrm{Mal}}$ | 1.0 | mM | Literature consensus |
| $K_m^{\mathrm{OAA,MDH}}$ | 0.040 | mM | Literature consensus |
| $[\mathrm{Mal}]$ | 0.30 | mM | Erecińska et al. (1988) |
| $V_{\mathrm{max}}^{\mathrm{CS}}$ | 14.5 | mM/min | Calibrated to place the extension in the same low-OAA regime as the baseline model |
| $K_m^{\mathrm{OAA,CS}}$ | 0.005 | mM | Literature consensus |
| $K_m^{\mathrm{AcCoA,CS}}$ | 0.010 | mM | Literature consensus |

**Table S15:** *Comparison of the clamped baseline and the dynamic-OAA sensitivity extension. The ASPNAT-off branch required stiff-solver fallback after LSODA failed the residual criterion, so the dynamic row is retained as a solver-checked sensitivity result rather than as an independent physiological steady state. Absolute flux magnitudes are not meant to be compared directly across the two OAA regimes.*

| Model | Steady-state OAA | $\mathbf{v}_{\mathrm{AAT}}$ on | $\mathbf{v}_{\mathrm{AAT}}$ off | Relief (%) |
|---|---|---|---|---|
| Reference clamped OAA | 160 nM | 0.1845 | 0.1410 | 30.9 |
| Dynamic OAA sensitivity state | about 0.3 nM | 0.0047 | 0.0035 | 34.0 |

**Table S16:** *Two-compartment stress-state rerouting sign test. Mitochondrial ASPNAT is the default biological premise; the external branch asks whether pathological rerouting of effective aspartate consumption would reverse the sign of AAT-linked support.*

| ASPNAT location | $\mathbf{v}_{\mathrm{AAT}}$ on | $\mathbf{v}_{\mathrm{AAT}}$ off | Relief (%) | $[\mathrm{Asp}]_{\mathbf{m}}$ on |
|---|---|---|---|---|
| Mitochondrial | 0.1351 | 0.0881 | +53.32 | 69.6 µM |
| External stress-rerouting branch | 0.0724 | 0.0881 | −17.80 | 72.4 µM |

**Table S17:** *Effective rate constants and the literature-scale fluxes they imply at baseline steady state.*

| Parameter | Value | Implied flux | Literature range |
|---|---|---|---|
| $k_{\text{export}}$ | 1.0 $\text{min}^{-1}$ | $J_{\text{export}} = 0.105$ mM/min | about 0.1 to 0.3 mM/min for AGC1-linked transport |
| $k_{\alpha\text{KG}}$ | 2.0 $\text{min}^{-1}$ | $J_{\alpha\text{KGDH}} = 0.185$ mM/min | about 0.2 to 0.4 mM/min for downstream oxidation |
| $k_{\text{NAA}}$ | 0.01 $\text{min}^{-1}$ | $J_{\text{NAA}} = 0.079$ mM/min | matrix removal with $\tau = 100$ $\text{min}^{\dagger}$ |

$^{\dagger}$This parameter describes matrix export on the timescale of the reduced model rather than whole-brain NAA turnover (I.-Y. Choi and Gruetter 2004).

**Table S 18:** *AGC1 flux implied by the first-order export model at representative matrix aspartate levels.*

| [Asp] ($\mu$M) | $\mathbf{J}_{\text{export}}$ (mM/min) |
|---|---|
| 50 | 0.050 |
| 100 | 0.100 |
| 150 | 0.150 |
| 200 | 0.200 |
| 500 | 0.500 |

**Table S19:** *Parameter confidence classes and plausible ranges used in the global uncertainty program.*

| Parameter | Baseline | LHS range | Class$^{\dagger}$ | Range basis |
|---|---|---|---|---|
| $K_m$ OAA | 50 $\mu$M | 30 to 100 $\mu$M | Measurement | Range across reported assays |
| $K_{\text{eq}}$ | 6.6 | 5.0 to 8.0 | Measurement | Thermodynamic measurements |
| $\text{IC}_{50}$ NAA | 0.85 mM | 0.5 to 1.5 mM | Measurement | Assay variability |
| $\text{IC}_{50}$ CoA | 0.42 mM | 0.2 to 0.8 mM | Measurement | Assay variability |
| [OAA] | 160 nM model reference | 50 to 500 nM | Physiology | From metabolic stress to mild activation |
| [AcCoA] | 14 $\mu$M | 5 to 50 $\mu$M | Physiology | From stress to carbon-replete states |
| $k_{\text{export}}$ | 1.0 $\text{min}^{-1}$ | 0.3 to 3.0 | Structural | Inferred from flux scale |
| $k_{\alpha\text{KG}}$ | 2.0 $\text{min}^{-1}$ | 1.0 to 4.0 | Structural | Tissue-dependent downstream oxidation |

| Parameter | Baseline | LHS range | Class† | Range basis |
|---|---|---|---|---|
| $V_{\mathrm{ASPNAT}}$ | 29.0 | 5 to 100 | Structural | Plausible effective capacity range |
| $k_{\mathrm{NAA}}$ | 0.01 $\mathrm{min}^{-1}$ | 0.005 to 0.02 | Structural | Plausible matrix removal range |

†Measurement-based values are anchored in direct assays. Physiology-derived values span plausible metabolic states. Structural values describe effective tissue-level parameters that are not directly measurable in vivo.

#### 2.4 Substrate-supply and thermodynamic-leverage decoupling

To separate substrate supply from thermodynamic leverage, we varied oxaloacetate and acetyl-CoA independently while keeping the rest of the reduced mitochondrial node unchanged. This analysis was designed as a qualitative node-level comparison, not as a quantitative reproduction of the intact-mitochondria efflux rates measured by Patel and Clark (1979). In the model, increasing acetyl-CoA increased absolute ASPNAT flux and lowered residual aspartate export, consistent with the pyruvate-supported reciprocal NAA/aspartate efflux behavior reported by Patel and Clark. By contrast, lowering oxaloacetate increased the relative thermodynamic leverage of the ASPNAT-mediated sink on forward AAT flux. Notably, even though the absolute rate of ASPNAT flux is smaller under low-oxaloacetate conditions, its proportional rescue of the near-equilibrium AAT reaction is much higher. The two dependencies therefore occupy different axes: acetyl-CoA supplies the fuel for absolute NAA synthesis, whereas low oxaloacetate defines the thermodynamic state in which product removal has greatest relative leverage.

**Table S20:** *Representative points from the OAA x acetyl-CoA decoupling sweep. ASPNAT flux is reported in the model's internal flux units. Residual aspartate export is the ASPNAT-on export flux expressed as a percentage of the matched ASPNAT-off export flux at the same boundary condition.*

| OAA (nM) | Acetyl-CoA ($\mu$M) | ASPNAT flux (model units) | Residual Asp export (% OFF) | Relative AAT relief (%) |
|---|---|---|---|---|
| 10 | 1 | 0.0097 | 84.4 | 14.3 |
| 10 | 14 | 0.0328 | 55.1 | 55.9 |
| 10 | 100 | 0.0515 | 38.3 | 96.6 |
| 160 | 1 | 0.0229 | 91.9 | 8.2 |
| 160 | 14 | 0.0792 | 74.8 | 30.9 |
| 160 | 100 | 0.1339 | 61.4 | 56.4 |
| 10000 | 1 | 0.0468 | 97.9 | 2.1 |
| 10000 | 14 | 0.1665 | 92.9 | 7.5 |
| 10000 | 100 | 0.3005 | 87.5 | 13.9 |

### 3 Supplementary Figures

These supplementary figures are restricted to computational safety checks that support the main narrative without repeating any current figure. They document the dynamic-OAA sensitivity extension, transport-law substitutions, and parameter-ensemble behavior used to test whether the sign and interpretation of the relief effect survive beyond the baseline presentation.

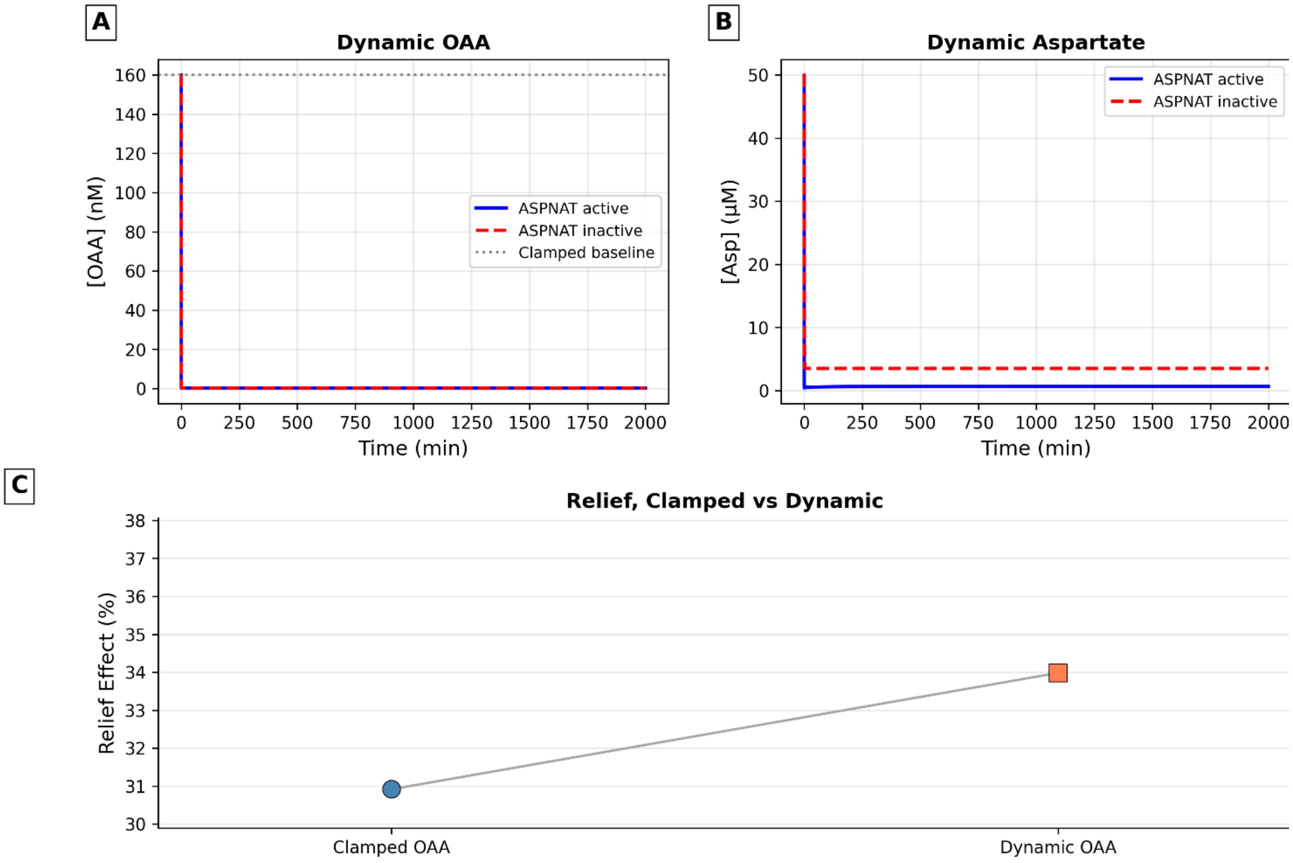


***Figure S1:*** *Dynamic-OAA sensitivity extension. Releasing the fixed-OAA boundary with generic thermodynamic malate dehydrogenase and citrate synthase terms preserved the positive direction of ASPNAT-mediated AAT relief after stiff-solver verification, but the absolute flux scale of this isolated extension remains subphysiological and should not be read as a neuronal OAA-calibrated whole-mitochondrion model. This figure therefore supports directional robustness of the relief mechanism, not a standalone physiological steady-state estimate.* ***A)*** *OAA trajectories show the dynamic boundary relaxing away from a fixed clamp.* ***B)*** *Aspartate trajectories retain the expected separation between ASPNAT-active and ASPNAT-inactive conditions.* ***C)*** *The dynamic-OAA simulation retained 33.98% relative AAT relief under the manuscript's canonical relief metric.*

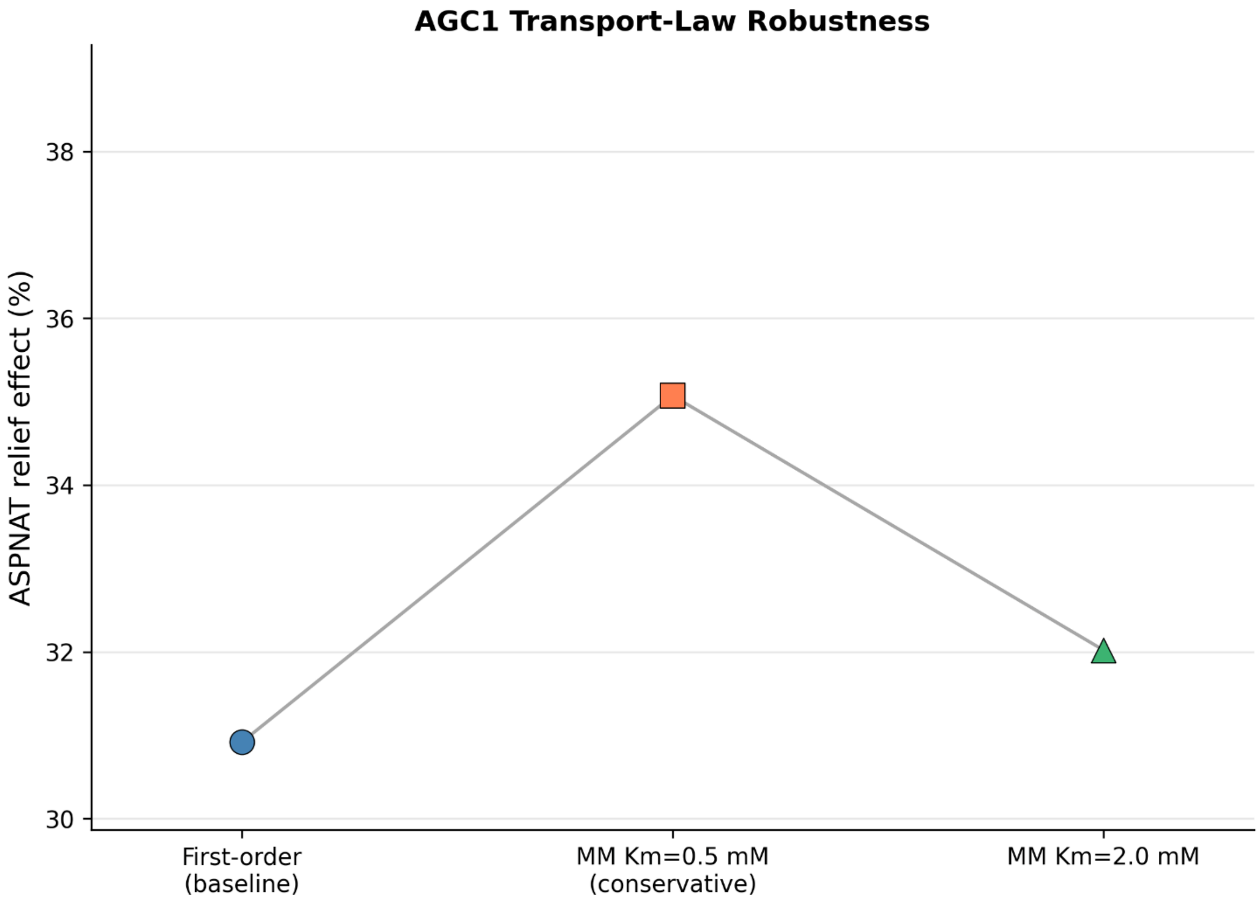


**Figure S2:** *AGC1 transport-law substitution check. Replacing the first-order export approximation with saturable Michaelis-Menten AGC1 kinetics preserves the positive sign of the relief effect across both tested parameterizations, yielding 35.1% relief at* $K_m = 0.5 mM$ *and 32.0% relief at* $K_m = 2.0 mM$*. This panel therefore functions as a transport-law robustness check rather than as a separate biological claim.*

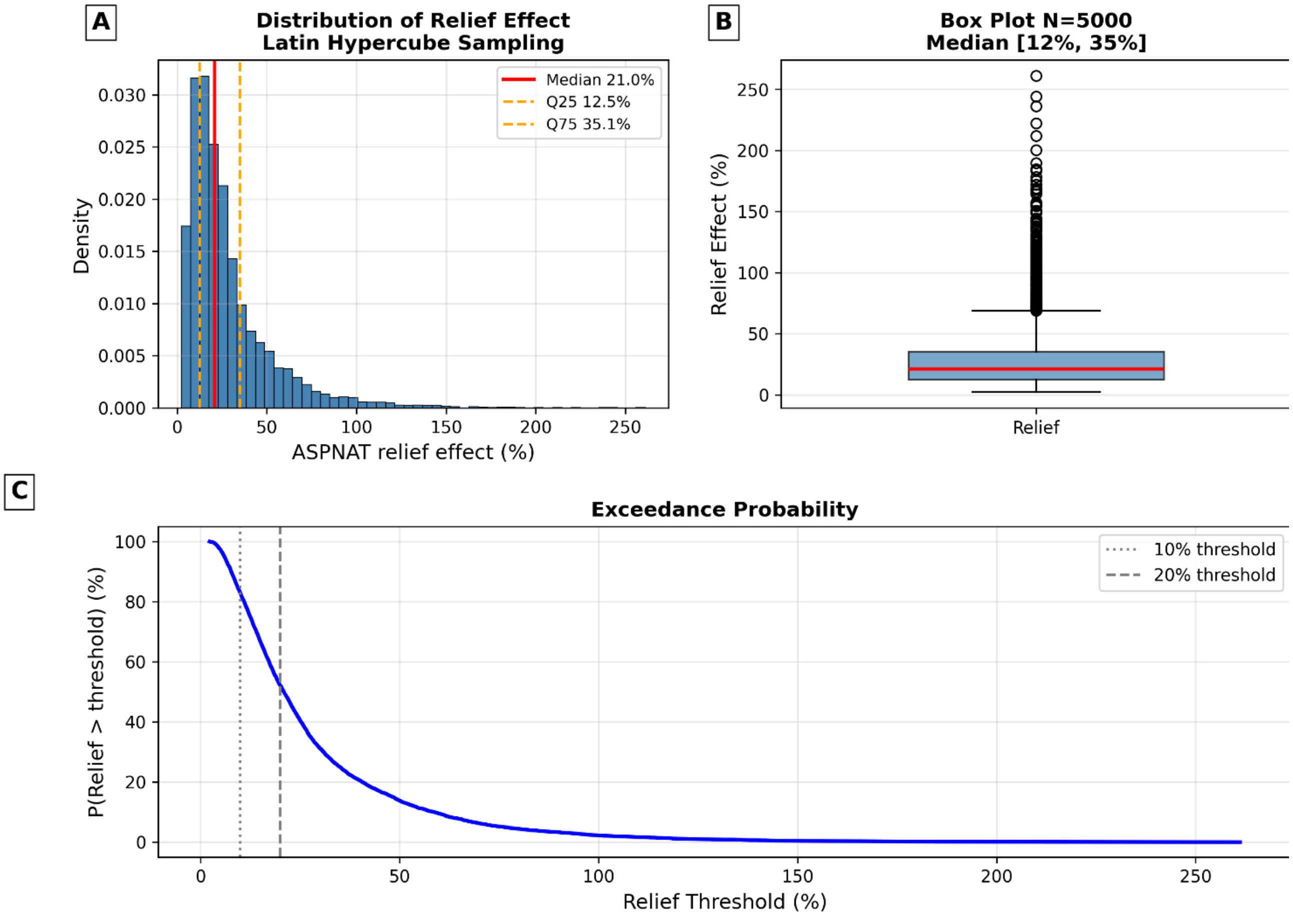


**Figure S3:** *Global uncertainty ensemble for the relief effect. All 5000 Latin Hypercube samples converged under the manuscript steady-state criterion, none produced negative relief, and the resulting distribution had a median relief of 20.95% with an interquartile range of 12.50 to 35.05%.* ***A*** *The density plot shows the distribution and quartiles.* ***B*** *The box plot summarizes the same ensemble.* ***C*** *The exceedance curve shows how robustness depends on the relief threshold.*

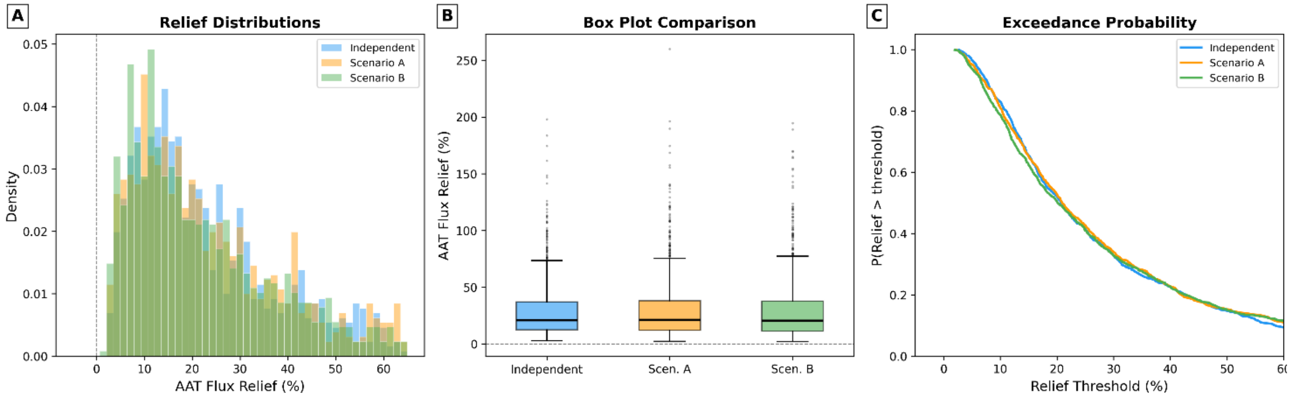


**Figure S4:** *Correlated-parameter robustness. Imposing strong covariance among OAA, acetyl-CoA, and export-related terms leaves the positive branch intact across all three tested ensembles.* ***A*** *Distribution overlays compare independent sampling with the two covariance scenarios.* ***B*** *Box plots*

*show similar central tendencies across scenarios.* ***C*** *Exceedance curves show that the qualitative positive-relief result is not a consequence of independent sampling alone.*


Alon, Uri. 2006. *An Introduction to Systems Biology: Design Principles of Biological Circuits*. Chapman; Hall/CRC. https://doi.org/10.1201/9781420011432.

Choi, In-Young, and Rolf Gruetter. 2004. "Dynamic or Inert Metabolism? Turnover of N-Acetyl Aspartate and Glutathione from D-[1-13C]glucose in the Rat Brain in Vivo." *Journal of Neurochemistry* 91 (4): 778–87. https://doi.org/10.1111/j.1471-4159.2004.02716.x.

Choi, Kiri, J. Kyle Medley, Matthias König, Kaylene Stocking, Lucian Smith, Stanley Gu, and Herbert M. Sauro. 2018. "Tellurium: An Extensible Python-Based Modeling Environment for Systems and Synthetic Biology." *Biosystems* 171: 74–79. https://doi.org/10.1016/j.biosystems.2018.07.006.

Dienel, Gerald A. 2019. "Brain Glucose Metabolism: Integration of Energetics with Function." *Physiological Reviews* 99 (1): 949–1045. https://doi.org/10.1152/physrev.00062.2017.

Erecińska, Maria, Malgorzata M. Zaleska, Itzhak Nissim, David Nelson, Fiorenzo Dagani, and Marc Yudkoff. 1988. "Glucose and Synaptosomal Glutamate Metabolism: Studies with [15N]Glutamate." *Journal of Neurochemistry* 51 (3): 892–902. https://doi.org/10.1111/j.1471-4159.1988.tb01826.x.

Hucka, Michael, Andrew Finney, Herbert M. Sauro, Hamid Bolouri, John C. Doyle, Hiroaki Kitano, Adam P. Arkin, et al. 2003. "The Systems Biology Markup Language (SBML): A Medium for Representation and Exchange of Biochemical Network Models." *Bioinformatics* 19 (4): 524–31. https://doi.org/10.1093/bioinformatics/btg015.

Jamshidi, Neema, and Bernhard O. Palsson. 2010. "Mass Action Stoichiometric Simulation Models: Incorporating Kinetics and Regulation into Stoichiometric Models." *Biophysical Journal* 98 (2): 175–85. https://doi.org/10.1016/j.bpj.2009.09.064.

Kimmich, George A., James A. Roussie, and Joan Randles. 2002. "Aspartate Aminotransferase Isotope Exchange Reactions: Implications for Glutamate/Glutamine Shuttle Hypothesis." *American Journal of Physiology-Cell Physiology* 282 (6): C1404–13. https://doi.org/10.1152/ajpcell.00487.2001.

Madhavarao, C. N., C. Chinopoulos, K. Chandrasekaran, and M. A. A. Namboodiri. 2003. "Characterization of the *N* -Acetylaspartate Biosynthetic Enzyme from Rat Brain." *Journal of Neurochemistry* 86 (4): 824–35. https://doi.org/10.1046/j.1471-4159.2003.01905.x.

McKenna, Mary C., Irene B. Hopkins, Steven L. Lindauer, and Penelope Bamford. 2006. "Aspartate Aminotransferase in Synaptic and Nonsynaptic Mitochondria: Differential Effect of Compounds That Influence Transient Hetero-Enzyme Complex (Metabolon) Formation." *Neurochemistry International* 48 (6–7): 629–36. https://doi.org/10.1016/j.neuint.2005.11.018.

Medley, J. Kyle, Kiri Choi, Matthias König, Lucian Smith, Stanley Gu, Joseph Hellerstein, Stuart C. Sealfon, and Herbert M. Sauro. 2018. "Tellurium Notebooks: An Environment for Reproducible Dynamical Modeling in Systems Biology." *PLOS Computational Biology* 14 (6): e1006220. https://doi.org/10.1371/journal.pcbi.1006220.

Moffett, John R., Brian Ross, Peethambaran Arun, Chikkathur N. Madhavarao, and Aryan M. A. Namboodiri. 2007. "N-Acetylaspartate in the CNS: From Neurodiagnostics to Neurobiology." *Progress in Neurobiology* 81 (2): 89–131. https://doi.org/10.1016/j.pneurobio.2006.12.003.

Moffett, John R., Suzannah B. Tieman, Daniel R. Weinberger, Joseph T. Coyle, and Aryan M. A. Namboodiri, eds. 2006. *N-Acetylaspartate: A Unique Neuronal Molecule in the Central Nervous System*. Vol. 576. Advances in Experimental Medicine and Biology. Boston, MA: Springer US. https://doi.org/10.1007/0-387-30172-0.

Moreno, Antonio, Brian D. Ross, and Stefan Blüml. 2001. "Direct Determination of the N-Acetyl-L-Aspartate Synthesis Rate in the Human Brain by 13C MRS and [1-13C]glucose Infusion." *Journal of Neurochemistry* 77 (1): 347–50. https://doi.org/10.1046/j.1471-4159.2001.t01-1-00282.x.

Palsson, Bernhard O. 2006. *Systems Biology: Properties of Reconstructed Networks*. Cambridge University Press. https://doi.org/10.1017/CBO9780511790515.

Patel, Tarun B., and John B. Clark. 1979. "Synthesis of N-Acetyl-L-Aspartate by Rat Brain Mitochondria and Its Involvement in Mitochondrial/Cytosolic Carbon Transport." *Biochemical Journal* 184 (3): 539–46. https://doi.org/10.1042/bj1840539.

Petzold, Linda R. 1983. "Automatic Selection of Methods for Solving Stiff and Nonstiff Systems of Ordinary Differential Equations." *SIAM Journal on Scientific and Statistical Computing* 4 (1): 136–48. https://doi.org/10.1137/0904010.

Porubsky, Veronica L., and Herbert M. Sauro. 2023. "A Practical Guide to Reproducible Modeling for Biochemical Networks." In *Computational Modeling of Signaling Networks*, edited by Lan K. Nguyen, 2634:107–38. Methods in Molecular Biology. New York, NY: Humana. https://doi.org/10.1007/978-1-0716-3008-2_5.

Ronowska, Anna, Andrzej Szutowicz, Hanna Bielarczyk, Sylwia Gul-Hinc, Joanna Klimaszewska-Łata, Aleksandra Dyś, Marlena Zyśk, and Agnieszka Jankowska-Kulawy. 2018. "The Regulatory Effects of Acetyl-CoA Distribution in the Healthy and Diseased Brain." *Frontiers in Cellular Neuroscience* 12: 169. https://doi.org/10.3389/fncel.2018.00169.

Sauro, Herbert M., and Frank T. Bergmann. 2010. "Software Tools for Systems Biology." In *Systems Biomedicine*, 289–314. Elsevier. https://doi.org/10.1016/B978-0-12-372550-9.00012-2.

Sauro, Herbert M., and Boris N. Kholodenko. 2004. "Quantitative Analysis of Signaling Networks." *Progress in Biophysics and Molecular Biology* 86 (1): 5–43. https://doi.org/10.1016/j.pbiomolbio.2004.03.002.

Smith, Lucian P., Frank T. Bergmann, Deepak Chandran, and Herbert M. Sauro. 2009. "Antimony: A Modular Model Definition Language." *Bioinformatics* 25 (18): 2452–54. https://doi.org/10.1093/bioinformatics/btp401.

Truckenmiller, M. E., M. A. A. Namboodiri, M. J. Brownstein, and J. H. Neale. 1985. "N-Acetylation of L-Aspartate in the Nervous System: Differential Distribution of a Specific Enzyme." *Journal of Neurochemistry* 45 (5): 1658–62. https://doi.org/10.1111/j.1471-4159.1985.tb07240.x.

Wang, Qinzhe, Mojun Zhao, Gwenn G. Parungao, and Ronald E. Viola. 2016. "Purification and Characterization of Aspartate N-acetyltransferase: A Critical Enzyme in Brain Metabolism." *Protein Expression and Purification* 119: 11–18. https://doi.org/10.1016/j.pep.2015.11.001.
